# Genomic, Functional and Structural Analyses Reveal Mechanisms of Evolutionary Innovation within the Sea Anemone 8 Toxin Family

**DOI:** 10.1101/2022.12.08.518931

**Authors:** Lauren M. Ashwood, Khaled A. Elnahriry, Zachary K. Stewart, Thomas Shafee, Muhammad Umair Naseem, Tibor G. Szanto, Chloé A. van der Burg, Hayden L. Smith, Joachim M. Surm, Eivind A.B. Undheim, Bruno Madio, Brett R. Hamilton, Shaodong Guo, Dorothy C.C. Wai, Victoria L. Coyne, Matthew J. Phillips, Kevin J. Dudley, David A. Hurwood, Gyorgy Panyi, Glenn F. King, Ana Pavasovic, Raymond S. Norton, Peter J. Prentis

**Affiliations:** School of Biology and Environmental Science, Faculty of Science, Queensland University of Technology, Brisbane, QLD 4000, Australia; Cancer Program, QIMR Berghofer Medical Research Institute, Brisbane, QLD, 4006, Australia; Medicinal Chemistry, Monash Institute of Pharmaceutical Sciences, Monash University, Parkville, VIC 3052, Australia; Centre for Agriculture and the Bioeconomy, Queensland University of Technology, Brisbane, QLD 4000, Australia; Department of Animal Plant & Soil Sciences, La Trobe University, Melbourne; Swinburne University of Technology, Melbourne, Victoria, Australia; Department of Biophysics and Cell biology, Faculty of Medicine, University of Debrecen, 4032, Debrecen, Hungary; Department of Anatomy, School of Biomedical Sciences, University of Otago, Dunedin, 9016, New Zealand; Department of Ecology, Evolution and Behavior, Alexander Silberman Institute of Life Sciences, The Hebrew University of Jerusalem, 9190401, Jerusalem, Israel; Centre for Ecological and Evolutionary Synthesis, Department of Biosciences, University of Oslo, PO Box 1066 Blindern, 0316 Oslo, Norway; Centre for Advanced Imaging, The University of Queensland, St Lucia, QLD 4072, Australia; Institute for Molecular Bioscience, The University of Queensland, St Lucia, QLD 4072, Australia; Centre for Microscopy and Microanalysis, The University of Queensland, St Lucia, QLD 4072, Australia; Research Infrastructure, Central Analytical Research Facilities, Queensland University of Technology, Brisbane 4000, QLD, Australia; ARC Centre for Innovations in Peptide and Protein Science, The University of Queensland, St Lucia, QLD 4072, Australia; School of Biomedical Sciences, Faculty of Health, Queensland University of Technology, Brisbane, QLD 4000, Australia; ARC Centre for Fragment-Based Design, Monash University, Parkville, Victoria 3052, Australia

**Keywords:** Sea anemone, toxin evolution;genome, neofunctionalization, peptide synthesis, disulfide connectivity

## Abstract

ShK from *Stichodactyla helianthus* has established the therapeutic potential of sea anemone venom peptides, but many lineage-specific toxin families in actinarians remain uncharacterised. One such peptide family, sea anemone 8 (SA8), is present in all five sea anemone superfamilies. We explored the genomic arrangement and evolution of the SA8 gene family in *Actinia tenebrosa* and *Telmatactis stephensoni*, characterised the expression patterns of SA8 sequences, and examined the structure and function of SA8 from the venom of *T*. *stephensoni*. We identified ten SA8 genes in two clusters and six SA8 genes in five clusters for *T. stephensoni* and *A. tenebrosa*, respectively. Nine SA8 *T. stephensoni* genes were found in a single cluster and an SA8 peptide encoded by an inverted SA8 gene from this cluster was recruited to venom. We show that SA8 genes in both species are expressed in a tissue-specific manner and the inverted SA8 gene has a unique tissue distribution. While functional activity of the SA8 putative toxin encoded by the inverted gene was inconclusive, its tissue localisation is similar to toxins used for predator deterrence. We demonstrate that, although mature SA8 putative toxins have similar cysteine spacing to ShK, SA8 peptides are distinct from ShK peptides based on structure and disulfide connectivity. Our results provide the first demonstration that SA8 is a unique gene family in actiniarians, evolving through a variety of structural changes including tandem and proximal gene duplication and an inversion event that together allowed SA8 to be recruited into the venom of *T*. *stephensoni*.

## Introduction

Sea anemones (order Actiniaria) rely on venom for their survival, deploying this chemical weapon during encounters with prey, predators and competitors [1]. In contrast to other cnidarians, however, the venom of actiniarians is dominated by peptide neurotoxins [2, 3]. These neurotoxins confer substantial clinical utility and have been utilised as molecular probes in studies of the nervous system for more than 40 years [4]. Characterised sea anemone neurotoxins can be classified into one of 12 pharmacological toxin families, based on which receptors and ion channels they target [5–8]. Sea anemone neurotoxins, however, can also be classified according to their structural scaffold [9]. For example, the six sea anemone potassium channel toxin families are characterised by different structural scaffolds [9]. Furthermore, these structural scaffolds are not always associated with a single toxin family. The inhibitor cystine knot (ICK) fold is among the most widely recruited motifs in animal venoms [10, 11] and cysteine spacing characteristic of the ICK motif has been identified in both sea anemone type 5 potassium channel toxins and acid-sensing ion channel toxins [9, 12]. Similarly, the β-defensin fold is observed in multiple sea anemone neurotoxin families [9]. Thus, much of the pharmacological diversity of venoms, including those of sea anemones, is often accounted for by a relatively small number of peptide scaffolds. Reflecting their amenability to functional innovation, these scaffolds—sometimes referred to as *privileged scaffolds*—have been recruited to venoms on numerous occasions, including within the same lineage.

ShK represents another privileged scaffold, with 7,329 proteins containing 13,829 ShK-like (ShKT) domains spanning multiple kingdoms documented in the Simple Modular Architecture Research Tool (SMART) database [6, 13]. In actiniarians, however, the ShK fold has only been observed within toxins belonging to the type 1 potassium channel toxin family [6, 9]. This motif is named after ShK, a 35-residue toxin isolated from the sea anemone, *Stichodactyla helianthus*, which potently blocks K_V_1.3 as well as other voltage-gated potassium channels [14–18]. An analogue of ShK (dalazatide, Shk-186) has shown remarkable efficacy in in preclinical animal models of autoimmune disease [15, 19–21] and entered phase 1 clinical trials for the treatment of plaque psoriasis [22]. Because of the therapeutic potential of ShK, sea anemone proteins containing ShK-like domains have received substantial research attention [23–27], but not all ShK-like peptides block ion channels. More recently, it was reported that multiple toxin-like peptides with the cysteine spacing characteristic of ShK are localised to neurons rather than venom-related structures in the starlet sea anemone, *Nematostella vectensis* [28]. One of these neuropeptides was a homolog of U-actitoxin-Avd8e, a member of the sea anemone 8 (SA8) toxin family [28].

SA8 putative toxins were first identified computationally in *Anemonia viridis* [29] and have since been noted in several other studies [30–33]. The SA8 gene family appears to be restricted to the order Actiniaria and thus far it has been identified in 14 sea anemone species, spanning the Actinioidea, Metridioidea and Edwardsioidea superfamilies [33]. Despite this, peptides from the SA8 putative toxin family have not been functionally validated as toxins and the tertiary structure of this family has not been resolved. Given the recent discovery of the ShKT cysteine motif in a *N. vectensis* SA8-like peptide, as well as the ongoing recruitment of neuropeptides into sea anemone venom [28], SA8 putative toxins could potentially function as toxins and/or neuropeptides in different species. Here, we characterise the genomic arrangement, evolution and expression patterns of SA8 genes in two species, *Actinia tenebrosa* (superfamily Actinioidea) and *Telmatactis stephensoni* (superfamily Metridioidea) using a combined proteomic, transcriptomic and genomic approach. Furthermore, we investigated the structure and function of the first reported SA8 peptide recruited to the venom of *T*. *stephensoni*. Finally, we examined whether the SA8 gene family can be distinguished from the ShK gene family or whether it represents an extension of this gene family.

## Materials and Methods

### Genome sequencing

#### Short-read DNA sequencing

DNA was extracted from *A. tenebrosa* using the E.Z.N.A Mollusc DNA Kit (Omega Bio-Tek). Tentacle tissue was homogenised in a tube containing ML1 buffer (Omega Bio-Tek) and a stainless-steel ball bearing (Qiagen) using a Tissuelyser II (Qiagen), before the remaining steps were performed following the manufacturer’s protocol. To extract DNA from *T. stephensoni*, tentacle tissue was homogenised in liquid nitrogen with a mortar and pestle. The sample was then processed following the QIAGEN Genomic-tip procedure. DNA quantity was determined using a Qubit 4 fluorometer. Genomic DNA for both *A. tenebrosa* and *T. stephensoni* was sequenced using 150-bp paired-end chemistry on an Illumina HiSeq 2500 and X10, respectively.

### PacBio long-read sequencing

For *A. tenebrosa*, high molecular weight (HMW) DNA was extracted using two rounds of the method described for short read DNA-seq from the same individual. HMW DNA was extracted from *T. stephensoni* according to the 10X Genomics salting-out protocol for DNA extraction from single insects [34]. In order to extract HMW DNA, minor modifications were made to this protocol, including the use of wide-bore pipette tips, low bind tubes and a gentle bead clean-up. DNA molecular weight was assessed using PippinPulse pulsed-field gel electrophoresis, while purity and quantity were determined using a Qubit fluorometer. Samples of sufficient purity (260:280 ratio of 1.8–1.9 and 260:230 ratio of 1.6–2.0) and quantity were sequenced.

Five micrograms of genomic DNA from each species was prepared using needle shearing and the BluePippin (SageScience) size-selection system. The SMRTBell® Template Prep Kit 1.0 (PacBio) was then used to prepare 14-kilobase libraries. Sequencing was undertaken on four sequencing chips using a 16-h run time on a PacBio Sequel®.

#### Hi-C sequencing

Hi-C sequencing was performed to generate a chromosomal-level assembly for *A. tenebrosa.* Tentacle tissue from *A. tenebrosa* was crosslinked using 1% formaldehyde. Chromatin extraction, library preparation and sequencing were performed by Phase Genomics (Seattle, WA).

### Genome assembly and polishing

PacBio reads from *A. tenebrosa* and *T. stephensoni* were assembled using SMART *denovo* [35] and HGAP4 (Pacific Biosciences, SMART Link), respectively. Assemblies were polished using raw PacBio reads and a custom Arrow pipeline (https://github.com/zkstewart/Genome_analysis_scripts/tree/master/pipeline_scripts/polishing_pipeline_scripts/arrow). Four iterations of Arrow polishing were carried out for *A. tenebrosa*, while three iterations were conducted for *T. stephensoni*. Subsequently, both assemblies underwent two rounds of polishing with a custom Pilon pipeline (https://github.com/zkstewart/Genome_analysis_scripts/tree/master/pipeline_scripts/polishing_pipeline_scripts/pilon) using Illumina short reads. Indel errors were removed from the polished *A. tenebrosa* genome assembly with deGRIT (https://github.com/zkstewart/deGRIT).

Redundancy removal for *A. tenebrosa* was performed using the Purge Haplotigs pipeline [36]. This pipeline involved alignment of raw reads to the genome using minimap2 [37] and sorting using samtools [38]. Repeat regions were accounted for by providing Purge Haplotigs the results of our repeat masking (see section 3 “Repeat library generation”) converted to BED file format with a custom script. Program parameters were set by manual inspection of the histogram produced by the pipeline (low: 20, mid: 75, high: 170).

Finally, proximity-guided assembly was carried out on our *A. tenebrosa* assembly by Phase Genomics (Seattle, WA) using the Proximo platform. The quality of genome assemblies were assessed using BUSCO [39] and a custom script (https://github.com/zkstewart/Genome_analysis_scripts/blob/master/genome_stats.py).

### Repeat library generation

A custom repeat library (CRL) was generated using homology and structure-based prediction in addition to *de novo* repeat prediction. Miniature Inverted-repeat Terminal Elements (MITEs) were predicted using MITE-HUNTER v.11-2011[40] and detectMITE v.20170425 [41]. The output MITE models of both programs were clustered using CD-HIT v.4.6.4 [42] with the same parameters employed by the detectMITE program to produce a non-redundant set of MITE models (cd-hit-est -c 0.8 -s 0.8 -aL 0.99 -n 5). In addition, a structure-based prediction of long terminal repeat retrotransposons (LTR-RTs) was performed using LTRharvest (GT 1.5.10) [43] and LTR_FINDER v.1.06 [44] following the recommended protocol indicated by LTR_retriever commit 8180c24 [45] to identify canonical and non-canonical (i.e., non-TGCA motif) LTR-RTs. The MITE and LTR-RT libraries were used to mask the genome assembly using RepeatMasker open-4.0.7 [46] with settings ‘-e ncbi -nolow -no_is -norna’. After this homology and structure-based modelling, *de novo* repeat prediction was performed using RepeatModeler open-1.0.11 [47] with the masked genome as input.

All repeat models predicted by the aforementioned programs were then curated to remove models that were potentially part of genuine protein-coding genic regions. This process first involved the removal of any models that were confidently annotated by either LTR_retriever or by RepeatModeler (i.e., were not classified as “Unknown”) from consideration as these were assumed to be true positives. The remaining nucleotide repeat models were translated into six reading frames and were searched using HMMER 3.1b2 [48] for a list of domain models associated with transposable elements (TEs). This list was generated by adding Pfam [49] and NCBI CDD [50] domains to a list of domains identified by a previous study [51]. The added Pfam domains were based upon visual inspection of domain prediction results for putative transposable elements and comparison to the “Domain organisation” graphics provided by the Pfam website to find models that were not likely to appear in non-TEs. The CDD domains were part of the cl02808 RT_like superfamily. Any repeat models that obtained a TE-associated domain prediction were assumed to be true positives, and were removed from consideration.

To remove repeat models that may be part of genuine protein-coding genes, we generated a database of known genes. This database consisted of UniProtKB/Swiss-Prot proteins (version 10/26/17) [52] in conjunction with the gene models of *N. vectensis* (v.2.0) [53], *Exaipasia diaphana* (v.1.1) [54], *Acropora digitifera* (v.0.9) [55], and *Hydra vulgaris* (NCBI annotation release 102, June 8, 2015) [56]. Putative transposons were removed in this database by the same process detailed above using HMMER 3.1b2 and the list of TE-associated domains (i.e., any sequences with TE-associated domain hits were removed). Repeat models that remained after this curation process were removed from the initial CRL if they obtained a BLASTx result with E-value more significant than 1E^-2^ when submitted as a query against the gene model database. This process resulted in a high-quality CRL which was used to soft-mask the *A. tenebrosa* genome using RepeatMasker (-e ncbi -s -nolow -no_is -norna -xsmall) for subsequent gene model prediction. Scripts were produced to automate this process and are available from https://github.com/zkstewart/Genome_analysis_scripts/tree/master/repeat_pipeline_scripts.

### Gene model prediction

Gene model prediction was performed using two complementary approaches: transcriptome-based gene model creation and *ab initio* gene model prediction.

#### RNA-seq read quality control and mapping

RNA-seq reads generated from a previous studies involving *A. tenebrosa* [57] and *T. stephensoni* [58] were used to assist in gene model prediction. Raw reads were quality trimmed using Trimmomatic [59] with parameters "ILLUMINACLIP:$TRIMDIR/adapters/TruSeq3-PE.fa:2:30:10 SLIDINGWINDOW:4:5 LEADING:5 TRAILING:5 MINLEN:25" based on those used by the Trinity *de novo* assembler [60, 61]. Trimmed sequences were aligned to each genome file using STAR 2.5.4b (commit 5dbd58c) using the 2-pass procedure to assist in the identification of intron splice sites. The resulting SAM files were converted into sorted BAM files using samtools [38].

#### Transcriptome-based gene model prediction

Multiple transcriptome building programs were used to create a ‘master’ transcriptome file for subsequent curation using the EvidentialGene tr2aacds pipeline [62] following the parameters and methodology of Visser *et al.* [63]. This involved *de novo* transcriptome assembly using SOAPdenovo-Trans v.1.03[64] and Velvet/Oases [65, 66] with multiple k-mer lengths (23, 25, 31, 39, 47, 55, and 63 for both plus 71 for SOAPdenovo-Trans only) alongside Trinity v.2.5.1 [60]. We differed from Visser *et al.* [63] from here on. For Trinity, we modified the parameters of Visser *et al.* [63] to not specify min_contig_length and we set min_kmer_cov = 2 and SS_lib_type = RF. All resultant *de novo* assemblies had sequences shorter than length 350 bp removed with a custom script. Genome-guided transcriptomes were constructed for each genome assembly using Trinity and Scallop v.0.10.2 [67] with the results of the STAR spliced alignment. For Trinity we specified genome_guided_max_intron = 21,000 to reduce false positives. This number was based on preliminary analysis of an Illumina-based *A. tenebrosa* assembly [68] and the *Exaiptasia diaphana* genome for which, of 152,518 total introns, only 51 are > 21kb (0.03 %) in the v.1.1 genome .gff. Scallop was run with default settings except where library_type = first. No minimum size cut-off was enforced for genome-guided transcriptomes as we assumed any short predictions would likely be more accurate than those built from *de novo* assembly. All transcriptomes were then concatenated and subjected to the EvidentialGene tr2aacds pipeline which rendered a master transcriptome consisting of non-redundant genes including alternative isoforms.

PASA v.2.2.0 [69] (commit af03820) was used to align the master transcriptome against the assembled genomes (using BLAT v.35 [70] and GMAP [71]) and to generate gene model predictions using default recommended parameters (i.e., transcriptome files were ‘cleaned’ using the seqclean tool, and validate_alignments_in_db.dbi were provided arguments MIN_PERCENT_ALIGNED=75, MIN_AVG_PER_ID=95, and NUM_BP_PERFECT_SPLICE_BOUNDARY=0).

#### BRAKER1 *ab initio* gene model prediction

Gene models were predicted by BRAKER v.2.0.6 [72] using the repeat library soft-masked genome assembly (--softmasking) and the BAM file produced by STAR for use as ‘hints’ during Augustus v.3.2.3 [73] gene prediction.

#### Combined gene model prediction

EvidenceModeler [74] was used to combine transcriptome-based gene model predictions with BRAKER1-derived *ab initio* gene models with parameters set to favour transcriptome models above *ab initio* models (Augustus model weight = 1, Transcript-based model weight = 10). Following this, a custom pipeline to curate gene models was generated, which is available from (https://github.com/zkstewart/Genome_analysis_scripts/blob/master/gene_annotation_pipeline/gene_model_curation/processing_pipeline.sh). This pipeline finds additional gene copies missed by prior annotation steps, and removes spurious models derived from transposable elements and from ribosomal RNA.

### Phylogenetic and comparative genomic analysis

#### Genomic data resources

In addition to the two genomes reported herein, publicly available genomic resources for sea anemones were obtained for comparative genomic analyses. These included a chromosome-level *N. vectensis* genome (Nvec200) [75], the *E. diaphana* genome (GCF_001417965.1) [54], and the *Paraphelliactis xishaensis* genome (version 3) [76]. The annotation of *N. vectensis* was modified by manually annotating a putative sea anemone 8 gene into its genome prior to further analysis.

#### Gene family expansion and contraction

Gene families were predicted for the five sea anemone species using OrthoFinder (version 2.5.4) [77, 78]. Single copy genes were aligned with MAFFT (version 7.311) [79] and used as input to IQ-TREE 2 [80] with an estimated divergence date of 500 mya for *Actinia* and *Nematostella*. Gene family counts and our time-calibrated tree were used as input to CAFE (Computational Analysis of gene Family Evolution) 5 [81] to identify gene families with expansion/contracted predicted to a P-value threshold of 0.05 (fixed orthogroups).

#### Enrichment analysis

Gene ontologies (GOs) [82] and PFAMs [83] were predicted for each of the five sea anemone species’ representative gene models using custom scripts (https://github.com/zkstewart/Genome_analysis_scripts/tree/master/annotation_table). In short, Mmseqs2 [84] was used to query gene models against the UniRef90 database [85] and sequence matches had associated GO terms obtained from the id_mapping.tab file provided by the UniProt Knowledgebase [52]. PFAM 35.0 domain models were downloaded, and sequences were searched for these domains using HMMER3 [48]. Chi-squared tests were used to identify annotation terms enriched in expanded gene families; these tests were performed using a custom script available from https://github.com/zkstewart/Various_scripts/blob/master/DGE/enrichment/cafe_enrichment_analysis.py.

#### Phylogenetic analysis

Relationships among 71 sea anemone full-length sequences (alignment length 121 amino acid residues) from the SA8 gene family were inferred under maximum likelihood (ML) in IQ-TREE v2.1.3 [80]. The best fitting model of amino acid evolution was identified by ModelFinder within IQ-TREE as LG [86], with rate variation among sites accommodated by a proportion of invariant sites and a gamma distribution. This LG+I+G model was favoured by both BIC and corrected AIC (AICc). Default tree search parameters were employed within IQ-TREE and following reconstruction of the ML tree, clade support values were evaluated with 1000 ultrafast bootstrap samples.

### Analysis of genome collinearity and classification of gene duplications

To identify similar proteins in inter-genome and self-genome comparisons, homology of amino acid sequences between the two species and within a single genome was examined using BLASTp (1xE^-5^). These BLASTp alignments were leveraged by MCScanX (default parameters) to identify collinear blocks. Additionally, the duplicate_gene_classifier program from the MCScanX package was used to classify the origins of duplicate genes in the genomes of *A. tenebrosa* and *T. stephensoni*, independently.

### Expression analysis of SA8 genes and identification of toxin-like transcripts

To examine the expression patterns of SA8 genes, we made use of tissue-specific reads for *A. tenebrosa* [33] and *T. stephensoni* [58]. Transcriptome assembly and differential gene analysis were conducted using the Trinity short read *de novo* assembler v2.5.1 [87] and edgeR Bioconductor package v3.3.1 [88], respectively, as described in Ashwood *et al.* [58]. Similarly, Trinotate v3.1.1 was used for functional annotation as per Ashwood *et al*. [58].

Transcripts with homology to known animal toxins were identified from BLASTp searches against the UniProt database by filtering for transcripts annotated with the “toxin activity” GO term (GO:0090729), excluding bacterial toxins. Consequently, weighted gene co-expression network analysis (WGCNA) one-step network construction and module detection (power = 7, mergeCutHeight = 0.25, minModuleSize = 10, deepSplit = 3) was used to identify correlation networks of toxin-like transcripts present in *T. stephensoni* and *A. tenebrosa* [89].

### Milked venom peptide identification and mass spectrometry imaging

Peptides present in the milked venom of *T. stephensoni* and *A. tenebrosa* were previously identified by Ashwood *et al*. [58] and Surm *et al*. [33], respectively. Using these signal peptide-containing milked venom peptides as queries, homology to transcriptomic sequence and ToxProt entries was determined using tBLASTn and BLASTp, respectively. Significant BLAST hits were defined by a threshold e-value of 1xE^-5^. Similarly, the mass spectrometry imaging (MSI) data for *T. stephensoni* utilised in the current study were generated as detailed previously [58]. The spatial distribution of peptides of interest was determined from the MSI dataset using SciLS Lab.

### Chemical/sequence space analysis

A database of 102 SA8 peptides was generated using SA8 sequences identified from 14 actiniarian species spanning three superfamilies [33], as well as the SA8 translated ORFs of *T. stephensoni* from the current study,. We analysed these SA8 sequences by position-specific biophysical property distance analyses, or sequence-space analyses [90]. Briefly, the multiple sequence alignment was converted to a matrix based on the physiochemical properties of the amino acids at each position in the alignment (gaps assigned column average values for each property), before dimensionality reduction (by principal component analysis) and group assignment (by Bayesian model based clustering) [90]. Additionally, SA8 sequences were aligned against ShK-like proteins (PF01549) and the homologous C-terminal regions of cysteine-rich secretory proteins (CRISPs; PF08562) available in the Pfam database [49], allowing SA8 putative toxins to be visualised in a sequence space with ShKT-domain containing sequences. Data were visualised with custom [R] scripts based on rgl [91]. WGCNA modules were mapped to the clusters identified in the sequence space of mature SA8 peptides, and correlation between the two categorical variables was assessed using the GoodmanKruskal package [92].

### Peptide synthesis and structure characterisation

#### Design of vector used for expressing SA8

A nucleotide sequence encoding the mature peptide sequence of SA8, containing codons optimised for expression in *Escherichia coli*, was synthesised by GenScript (Piscataway, NJ, USA). The sequence was subcloned into a modified pET32a vector (Novagen, Madison, WI, USA) using *Bam*HI and *Eco*R1 restriction sites (**Figure 1**). This modified pET32a vector encodes a MalE signal sequence (for targeting the fusion protein to the *E. coli* periplasm), a polyhistidine (His_6_) affinity tag (for nickel affinity chromatography used for peptide purification), a maltose binding protein (MBP) fusion tag (to enhance the solubility of the expressed peptide), and a tobacco etch virus (TEV) protease recognition site (ENLYFQ) immediately preceding the coding region for SA8.

**Fig. 1.**
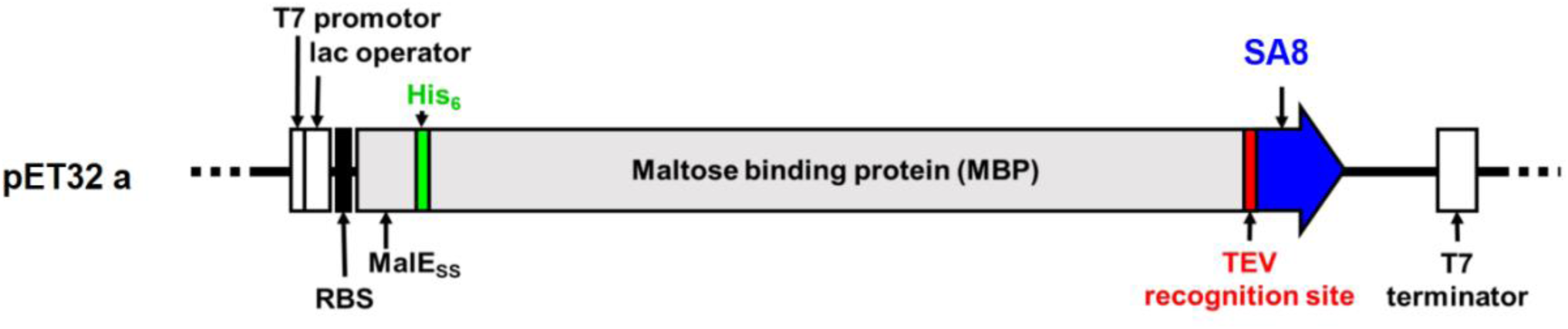
Schematic of pET32a expression vector used for expression of SA8 in the periplasm of *E. coli*. Abbreviations: RBS= ribosome binding site; MalESS= MalE signal sequence; His_6_= poly-histidine affinity tag; TEV= TEV protease recognition site; SA8= sea anemone 8

#### Recombinant expression of SA8

The pET32a-SA8 plasmid was transformed into *E. coli* strain BL21 (DE3) using a standard heat shock protocol [93]. Single colonies were inoculated into 5 mL culture medium (Luria-Bertani (LB) plus 100 μg/mL of ampicillin) and grown overnight at 37 °C with shaking at 150 rpm. The overnight starter culture was used to inoculate LB medium containing 100 μg/mL of ampicillin, then the culture was grown until it attained an optical density at 600 nm (OD_600_) of 0.6–0.8. Isopropyl β-D-1-thiogalactopyranoside (IPTG) was added to a final concentration of 0.5 mM to induce expression, and the culture was grown overnight at room temperature with shaking at 180 rpm. Induced cells were harvested by centrifugation for 15 min at 2,408 *g* and 4 °C, and the cell pellets were stored at –80 °C.

#### Peptide purification

The His_6_-MBP-SA8 fusion protein was recovered from the bacterial periplasm by osmotic shock [94]. Cell pellets were thawed on ice to avoid lysis, resuspended in 30 mL 100 mM Tris, 30% sucrose, 2 mM EDTA pH 8.0, and incubated for 10 min at 4 °C. The suspension was centrifuged for 15 min at 4 °C and 5,251 *g*. The pellets were resuspended in 30 mL of ice-cold water and 5 mM MgCl_2_, then incubated for 10 min at 4 °C before centrifugation for 15 min at 4 °C and 21,002 *g*. The supernatant containing soluble His_6_-MBP-SA8 fusion protein was diluted in 20 mM Tris, 150 mM NaCl, 5% glycerol pH 7.5 (TNG buffer) and incubated for 30 min with 5 mL Ni-nitrilotriacetic acid (Ni-NTA) Fast Flow resin (GE Healthcare, Uppsala, Sweden) in a gravity-fed column to capture the fusion protein via its affinity to Ni-NTA. The resin was washed with 25 mM imidazole to remove weakly bound proteins. The His_6_-MBP-SA8 fusion protein was then eluted with 200 mM imidazole, and then it was concentrated by centrifugal filtration (Amicon® Ultra, Millipore) and desalted using a PD-10 column (GE Healthcare, Pittsburgh, PA, USA) to remove imidazole. The fusion protein was incubated with His_6_-TEV protease (produced in-house; 40 μg per mg of fusion protein) for 16 h with shaking at 100 rpm and 30 °C. The cleavage mixture was then passed over Ni-NTA Fast Flow resin to remove His_6_-MBP and His_6_-TEV protease, and the eluate containing the liberated SA8 was collected for further purification using reversed-phase high performance liquid chromatography (RP-HPLC). RP-HPLC was performed on a Vydac C18 column (250 x 10 mm) using a flow rate of 1 mL/min and a gradient of 30%–50% solvent B (0.1% trifluoroacetic acid (TFA) in 90% acetonitrile) in solvent A (0.1% TFA in water) over 40 min with absorbance monitored at 214 nm. Fractions containing SA8 was collected and assessed by liquid chromatography-mass spectrometry (LC-MS). Fractions containing SA8 (>99%) were lyophilised for NMR studies, mapping disulfide connectivity, and functional assays.

#### NMR spectroscopy and disulfide connectivity

A SA8 sample for NMR experiments was prepared by dissolving lyophilised peptide in 90% H_2_O and 10% ^2^H_2_O and adjusting the pH by addition of 0.1 N HCl and 0.1 M NaOH. NMR spectra were acquired on a 600 MHz Bruker NMR spectrometer (Billerica, MA) equipped with a cryogenically cooled TCI probe. One-dimensional (1D) ^1^H NMR spectra were acquired over the temperature range 5–30 °C in increments of 5 °C to investigate the effects of temperature on the spectrum before selecting conditions for structure determination. Bruker TopSpin, USA, version 3.6.1, was used for processing all spectra.

The disulfide connectivity was determined by analysing pepsin-digested recombinant, oxidised SA8 peptide using liquid chromatography tandem mass spectrometry (LC-MS/MS). 5 µg SA8 was dissolved in 10 µL 10 mM hydrochloric acid and digested with 20 ng/µL pepsin (Merck, cat. No. 10108057001) at 37 °C for 6 h. The digested sample was diluted with formic acid (FA) to yield a final concentration of 0.3 ng/µL in 15 µL 0.5 % FA, from which 1 µL was analysed by LC-MS/MS. The sample was separated with a nanoElute nano-HPLC (Bruker, Germany) across a Bruker TEN C18 nano separation column (100 mm length, 75 µm inner diameter, 1.9 µm particle size, 120 Å pore size) heated to 50 °C, using a gradient of 5–30 % solvent B (90 % acetonitrile [ACN] in 0.1 % formic acid [FA]) in solvent A (0.1 % FA) over 18 min before a step gradient to 95 % B for 18 min at a flow rate of 0.5 µL/min. The eluting peptides were analysed with an in-line Bruker timsTOF Pro operated in positive, data-dependent analysis, parallel accumulation-serial fragmentation (DDA-PASEF) mode with a cycle time of 0.5 s and an m/z range of 350–2200. The resulting MS and MS2 spectra were manually compared to a list of all possible peptic fragment precursors and their fragment ions estimated from all theoretical disulfide permutations.

#### Co-elution of native and recombinant SA8

Crude extract containing the native SA8 peptide was isolated using a protocol based on that of Honma *et al.* [95]. Briefly, tissue from the body column of *T. stephensoni* was placed into tubes containing a single stainless-steel ball bearing (Qiagen) and homogenised using a Tissuelyser II (Qiagen). Subsequently, Milli-Q H_2_O was added to the tubes containing the macerate, which were then centrifuged for 15 min at 18,000 *g*, allowing the supernatant to be recovered. A second round of homogenisation and centrifugation was conducted after additional H_2_O was added to the tubes containing pelleted material. The supernatant from both rounds of extraction were combined, then flash frozen in liquid nitrogen and lyophilized.

The venom extract sample was diluted with 500 µL of Milli-Q water and centrifuged for 1 min at 17,680 *g*. The sample was loaded onto Sep-Pak C18 Vac cartridge/5000 mg (Waters) that was previously equilibrated with 1 mL methanol and 1 mL Milli-Q water. The sample was then washed with 500 µl of Milli-Q water and eluted with 750 µl of TFA/Milli-Q/acetonitrile (1:70:29). The eluate was evaporated under a constant stream of N_2_. The residue was reconstituted in Milli-Q water (80 µL) and analysed using LC-MS. An analytical Luna C8 LC column (100 mm x 2 mm) was utilised for co-elution of native and the recombinant peptides with a gradient of 0–60% buffer B (90% ACN, 0.1% trifluoroacetic acid [TFA]) in buffer A (0.1% TFA) and constant flow rate of 0.2 mL/min over 10 min.

#### *In vitro* refolding of reduced recombinant SA8 peptide

The recombinant SA8 peptide was reduced and unfolded in 10 mM dithiothreitol (DTT), 20 mM Tris at pH 8. Complete reduction occurred after approximately 30 min, as confirmed using LC-MS. Following reduction, a PD-10 column was used to desalt the sample and remove DTT. The reduced SA8 was then oxidised using 0.1 M NH_4_HCO_3_ buffer at pH 8, with complete oxidation occurring after 8 h, as verified using LC-MS. 1D ^1^H NMR spectra were acquired at pH 3.5 and 25°C for the major re-oxidised product to check folding.

### *In vitro* and *In vivo* functional testing

#### Patch clamp electrophysiology

Chinese hamster ovary (CHO) and human embryonic kidney (HEK 293) cells were grown in DMEM-high glucose supplemented with 10% FBS, 2 mM L-glutamine, 100 U/mL penicillin-g, and 100 μg/mL streptomycin (Invitrogen) at 37°C in a 5% CO_2_ and 95% air humidified atmosphere. Cells were passaged twice per week following a 5-min incubation in PBS containing 0.2 g/L EDTA (Invitrogen, Carlsbad, CA).

CHO cells were transiently transfected with plasmids encoding hK_V_1.1, hK_V_1.2, hK_V_10.1, hKCa3.1, hTRPA1, or hNa_V_1.7 using Lipofectamine 2000 (Invitrogen, Carlsbad, CA) following the manufacturer’s protocol, then cultured under standard conditions. The cells were co-transfected with a plasmid encoding green fluorescent protein (GFP) at a molar ratio of 10:1 except for hK_V_1.1, hK_V_1.2, and hKCa3.1 for which GFP-tagged ion channel vectors were used. Currents were recorded 24–36 h after transfection. GFP-positive transfectants were identified with a Nikon Eclipse TS100 fluorescence microscope (Nikon, Tokyo, Japan) using bandpass filters of 455–495 nm and 515–555 nm for excitation and emission, respectively, and these cells were used for current recordings (>70% success rate for transfection). HEK 293 cells stably expressing the hKv11.1 channel and mKCa1.1 channel were used. hTRPV1 channels were expressed in a stable manner in CHO cells. Transfected cells were washed twice with 2 mL of extracellular solution (ECS, see below) and re-plated onto 35-mm polystyrene cell culture dishes (Cellstar, Greiner Bio-One). hK_V_1.3 currents were recorded from activated peripheral blood mononuclear cells (PBMCs, see below) 3–4 days after activation. Human veinous blood was obtained from anonymised healthy donors. PBMCs were isolated using Histopaque1077 (Sigma-Aldrich Hungary, Budapest, Hungary) density gradient centrifugation. Cells obtained were resuspended in RPMI 1640 medium containing 10% fetal calf serum (Sigma-Aldrich Hungary, Budapest, Hungary), 100 μg/mL penicillin, 100 μg/mL streptomycin, and 2 mM L-glutamine, seeded in a 24-well culture plate at a density of 5–6 × 10^5^ cells/mL, and grown in a 5% CO_2_ incubator at 37°C for 3–5 days. Phytohemagglutinin A (PHA, Sigma-Aldrich Hungary, Budapest, Hungary) was added to the medium at 10 μg/mL to amplify the K_V_1.3 expression. Cells were washed gently twice with 2 mL of ECS for the patch-clamp experiments.

Conventional whole-cell patch-clamp electrophysiology [96] was used to record ionic currents. Micropipettes were pulled in four stages using a Flaming Brown automatic pipette puller (Sutter Instruments, San Rafael, CA) from Borosilicate Standard Wall with Filament aluminum-silicate glass (Harvard Apparatus Co., Holliston, MA) with tip diameters between 0.5 and 1 μm and heat polished to a tip resistance of 2–8 MΩ. All measurements were carried out by using Axopatch 200B amplifier connected to a personal computer using Axon Digidata 1550A data acquisition hardware (Molecular Devices, Sunnyvale, CA). In general, the holding potential was −120 mV. Records were discarded when leak at the holding potential was > 10% of peak current at the test potential. Series resistance compensation up to 70% was used to minimise voltage errors and achieve proper voltage-clamp conditions. Experiments were performed at room temperature (20–24°C). Data were analysed using Prism 8 (GraphPad, CA, USA) and ClampFit 10.5 software (Molecular Devices Inc., Sunnyvale, CA). Before analysis, whole-cell current traces were corrected for ohmic leakage and digitally filtered with a three-point boxcar smoothing filter. For hKCa3.1, hTRPA1 and hTRPV1 the reversal potential for K^+^ was determined and only those currents were analysed for which the reversal potential fell into the range of the theoretical reversal potential ± 5 mV (–86.5 ± 5 mV for KCa3.1, and 0 ± 5 mV for hTRPA1 and hTRPV1).

Solutions: All salts and components of the solutions were purchased from Sigma Aldrich, Budapest, Hungary. For hK_V_1.1, hK_V_1.2, hK_V_1.3, hK_V_10.1, mKCa1.1, and hNa_V_1.7 the ECS contained 145 mM NaCl, 5 mM KCl, 2.5 mM CaCl_2_, 1 mM MgCl_2_, 10 mM HEPES, and 5.5 mM glucose (pH 7.35 with NaOH), while the intracellular (pipette) solution (ICS) contained 140 mM KF, 2 mM MgCl_2_, 1 mM CaCl_2_, 11 mM EGTA, and 10 mM HEPES (pH 7.22 with KOH). For recordings of hNav1.7 currents, the composition of ICS was 105 mM CsF, 10 mM NaCl, 10 mM EGTA, and 10 mM HEPES, (pH 7.2 with CsOH). For recordings of hKv11.1 currents, the ECS contained 140 mM choline-chloride, 5 mM KCl, 2 mM MgCl_2_, 2 mM CaCl_2_, 0.1 mM CdCl_2_, 20 mM glucose, and 10 mM HEPES (pH 7.35 with NaOH) and the ICS contained 140 mM KCl, 10 mM EGTA, 2 mM MgCl_2_, and 10 mM HEPES (pH 7.3 with KOH). hKCa3.1 currents were recorded with an ECS of the following composition: 160 mM L-aspartic acid sodium salt, 5 mM KCl, 2.5 mM CaCl_2_, 1.0 mM MgCl_2_, 5,5 mM glucose and 10 mM HEPES (pH 7.4 with NaOH), while the composition of ICS was 150 mM L-aspartic acid potassium salt, 5 mM HEPES, 10 mM EGTA, 8.7 mM CaCl_2_, and 2 mM MgCl_2_ (pH 7.22 with KOH) giving ∼1.2 µM free Ca^2+^ to fully activate the KCa3.1 current [97]. For recordings of TRPA1 and TRPV1 currents, the ECS and ICS contained 150 mM NaCl, 2 mM Na-EDTA, and 10 mM HEPES (pH 7.35 with NaOH). hTRPA1 and hTRPV1 currents were activated by 100 µM allyl-isothiocyanate (AITC) and 1 µM capsaicin, respectively, and SA8 was added after complete activation of the current. SA8, activators and positive controls were dissolved in ECS, except for tetraethylammonium (TEA^+^) that was stored in 100 mM stock at 4°C. The osmolarity of the ECS and ICS were 302–308 mOsM and ∼ 295 mOsM, respectively. SA8 and positive controls were dissolved in ECS supplemented with 0.1 mg/mL bovine serum albumin (BSA; Sigma-Aldrich Hungary, Budapest, Hungary). Bath perfusion around the measured cell with different extracellular solutions was achieved using a gravity flow microperfusion system using a rate of 0.5 mL/min. Excess fluid was removed continuously.

Voltage protocols: For measurement of hK_V_1.1–1.3 and hK_V_10.1, currents, voltage steps to +50 mV were applied from a holding potential of −120 mV every 15 or 30 s and the peak current was measured. Channels were activated by series of depolarisation pulses to +50 mV from a holding potential of –120 mV. For K_V_1.1, 50-ms-long activating stimuli were used every 15 s. Due to the highly variable activation kinetics of K_V_1.2 [98] 200-ms-long pulses were applied every 15 s to maximise open probability of the channel. Due to the slow inactivation kinetics of K_V_1.2, the currents did not inactivate even at 200-ms-long depolarisation pulses. K_V_1.3 currents were evoked by 15-ms-long depolarisation pulses to +50 mV from –120 mV every 15 s. The use of such a short pulses every 15 s was sufficient to fully activate the channels and ensured that there was no cumulative inactivation of K_V_1.3. Positive controls were applied at a concentration equivalent to their *K*_d_ values (14 nM charybdotoxin (ChTx) for K_V_1.2, 0.3 mM TEA^+^ for K_V_1.1, 2µM Astemizole for K_V_10.1 and 10 mM TEA^+^ for K_V_1.3 and K_Ca_1.1). The approximate 50% reduction in the current amplitude in the presence of these compounds was an indicator of both the ion channel and the proper operation of the perfusion system. For mKCa1.1 channels, a voltage step to +100 mV from a holding potential of –100 mV was used. Pulses were delivered every 10 s. hKCa3.1 currents were elicited every 15 s with voltage ramps to +50 mV from a test potential of –120 mV at a rate of 0.85 mV/ms. The holding potential was set to –85 mV. For hK_V_11.1 channels, currents were evoked with a voltage step to +20 mV followed by a step to –40 mV, during which the peak current was measured. The holding potential was –80 mV, and pulses were delivered every 30 s. hTRPA1 and hTRPV1 currents were elicited every 5 s with 200-ms-long voltage ramps to +50 mV from a test potential of –50 mV. The cells were held at 0 mV during the subsequent pulses. The TRPA1 and TRPV1 currents were activated by 100 µM AITC and 1 µM capsaicin, respectively. Currents through hNa_V_1.7 channels were evoked every 15 s with a voltage step to +50 mV from a holding potential of –120 mV. The remaining current fraction (RCF) at a given molar concentration was calculated as *I/I_0_*, where *I_0_* is the peak current in the absence, and *I* is the peak current at equilibrium block or in the absence of inhibition after ∼2 min perfusion by SA8. Data points on concentration–response curves represent the mean of four individual measurements. These curves were fitted via simple linear regression according to the equation Y = 1 + slope × [toxin], where Y is the reciprocal of the remaining current fraction (1/RCF) and [toxin] is the molar concentration of SA8. The reciprocal of the slope yielded the IC_50_ value.

#### Injection activity assay in *Drosophila melanogaster*

The activity of the recombinant SA8 peptide in *Drosophila melanogaster* was assessed using an injection activity assay [99]. The mass of female *D. melanogaster* used for this experiment ranged from 0.8 to 1.1 mg. Five doses of SA8 (0.00005, 0.0005, 0.005, 0.05 and 0.5 µg per fly) were each administered to eight *D. melanogaster* (*n* total=40), with observations of paralysis and mortality made at 2 and 24 h post-injection. Dose-response data were analysed as described previously [99].

## Results and Discussion

### Genome assemblies of Actinia tenebrosa and Telmatactis stephensoni

The genome assembly for *A. tenebrosa* generated a 312-Mb assembly scaffolded into 18 pseudomolecules with a N50 of 16.5 Mb, with 38 contigs not assigned to scaffolds (**Table 1**). The genome assembly for *T. stephensoni* resulted in a 484-Mb assembly with a contig N50 of 602 Kb. Completeness scores (BUSCO) for *A. tenebrosa* and *T. stephensoni* assemblies are on par with other long-read actiniarian genomes [54, 75, 100, 101]. In contrast, contig N50 values for the current *A. tenebrosa* and *T. stephensoni* genomes are higher than both previously assembled short- and long-read genomes.

**Table 1.** Comparison of assembly metrics among actiniarian genomes. Abbreviations: TELST= T. stephensoni; ACTTE= *A. tenebrosa*; ACTTE*= draft *A. tenebrosa* [68]; ACTEQ= *Actinia equina* [100]; NEMVE= *N. vectensis* [101]; EXADI= *E. diaphana* [54]

| **Annotation metrics** | **TELST** | **ACTTE** | **ACTEQ** | **ACTTE\*** | **NEMVE** | **EXADI** |
| --- | --- | --- | --- | --- | --- | --- |
| Assembly size (Mbp) | 484 | 312 | 409 | 238 | 356 | 256 |
| Contig N50 (kbp) | 602 | 716 | 492 | 188 | 472 | 442 |
| Scaffold N50 (kbp) | - | 16,458 | - | - |  | - |
| Number of chromosomes | - | 18 | - | - | - | - |
| Percent repetitive DNA | 27.3 | 32.3 | - | 19.57 | 26 | 26 |
| BUSCO (%) | 93.8 | 93.5 | 94.0 | 89.6 | 91.6 | 87.3 |

Repeat content analysis of the *A. tenebrosa,* and *T. stephensoni* genomes found that 32.3% (101 Mb) and 27.3% (122 Mb), respectively, of these assemblies are repetitive. Similar proportions of repetitive DNA were observed in the *N. vectensis* and *E. diaphana* genome assemblies [54, 101]. LTR, MITEs and unknown repeats were the most observed repeat classes found in both sea anemone species, but the relative abundance of each class varied between species (**Supplementary Table 1**). LTRs were the most abundant repeat type found in the *A. tenebrosa* assembly, followed by MITEs and unknown repeats. In contrast, Surm *et al* [68] reported that MITES were the most common repeat type in the draft genome of *A. tenebrosa*, followed by unknown repeats. MITEs were the most abundant repeat class observed in the *T. stephensoni* assembly followed by LTRs and unknown repeats.

For *A. tenebrosa*, we predicted 53,560 protein coding genes (59,465 with isoforms included) with a BUSCO genome completeness score of 97.3%. We predicted 50,712 protein coding genes (54,933 with isoforms included) in *T. stephensoni* with a BUSCO genome completeness score of 96.6%. Based on these BUSCO scores, the completeness of our assemblies is comparable to that of the *A. equina* genome (97%) [100]. We predicted twice as many genes compared to previous sea anemone genome assemblies [54, 68, 75, 100, 101]. Furthermore, we predicted more protein coding genes in *A. tenebrosa* compared to the other assembly for this species as Surm *et al.*[68] produced a draft genome using short read sequences only. This resulted in a significant underestimation of gene number in the previous *A. tenebrosa* genome assembly.

### Expansion of gene families containing putative toxins within *T. stephensoni*

The phylogenetic tree generated for the five species is consistent with previous phylogenies [102], with *N. vectensis* sister to a clade containing *A. tenebrosa* and the three Metridioidean species. CAFE was then used to model the evolution of gene family sizes across five actiniarian species spanning the superfamilies Actinioidea, Edwardsioidea and Metridioidea (**Figure 2**). This analysis revealed *T. stephensoni* had the largest number of expansions and the fewest contractions of gene families. In fact, all three Metridioidean species had a greater number of gene family expansions compared to *A. tenebrosa* and *N. vectensis*. Gene families that encode putative toxins were common in the expanded gene families, with 21 expanded toxin gene families in *T. stephensoni* compared to 15, 10, 4 and 2 for *E. diaphana*, *P. xishaensis, A. tenebrosa* and *N. vectensis*, respectively (**Supplementary Table 2**). Orthogroups containing ShK-like, peptidase M12A, phospholipase A2 and other putative toxins were expanded in *T. stephensoni*, while orthogroups containing insulin-like growth factor binding protein (IGFBP) like and Factor V-like putative toxins were expanded in *A. tenebrosa.* Furthermore, CAFE results detected an expansion of the SA8 family only in *Telmatactis,* a pattern also observed by Surm *et al.* [33], with a contraction of this gene family occurring in *N. vectensis*. Thus, the SA8 family appears to have a unique expansion event in *Telmatactis* and the potential mechanisms for this expansion should be examined.

**Fig. 2.**
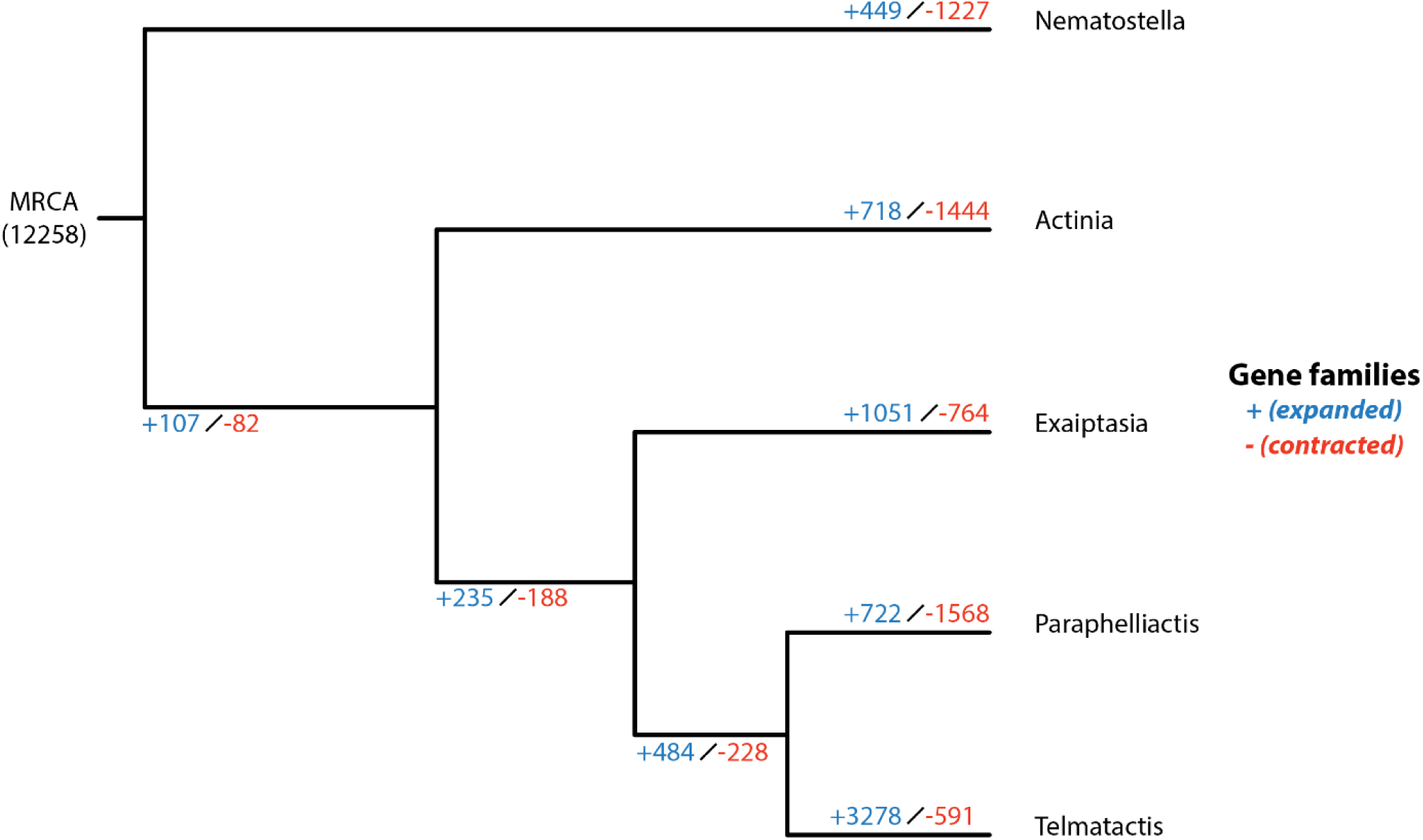
CAFE expansion/contraction tree highlights increased number of expanded gene families *in T. stephensoni* relative to other sea anemone species. Species: Nematostella = *N. vectensis*; Actinia = *A. tenebrosa*; Exaiptasia= *E. diaphana*; Paraphelliactis= *P. xishaensis*; Telmatactis *= T. stephensoni*

### Size and sequence divergence within the SA8 family coincides with phylogeny of actiniarian superfamilies

Multiple SA8 genes were identified in the genomes of *A. tenebrosa* and *T. stephensoni*, but only a single SA8 gene was identified and manually annotated in the most recent *N. vectensis* genome [75]. This single *N. vectensis* SA8 gene was comprised of three regular exons, ranging from 29 to >120 bp. The number of genes and their arrangement also differed significantly between *A*. *tenebrosa* and *T*. *stephensoni* (**Figure 3A**). In *T. stephensoni*, 10 SA8 genes were identified, nine of which were tandemly clustered on a single contig. In contrast, only six SA8 genes were identified in the *A. tenebrosa* genome on multiple pseudomolecules. Additionally, we noted distinct differences in the genomic arrangement of the SA8 gene family in relation to taxonomy. The six *A. tenebrosa* genes were located across four scaffolds, with multiple SA8 genes present on both scaffolds 1 and 11. The two SA8 genes on scaffold 11 were arranged tandemly, but in scaffold 1, the SA8 genes were 1.5 Mb apart and had different orientations (non-tandem). MCScanX analysis indicated that the cluster of nine SA8 genes in *T. stephensoni* originated from two episodes of tandem duplication involving eight genes collectively, with the remaining clustered SA8 gene originating from a proximal duplication event. The non-clustered *T. stephensoni* SA8 gene appears to have originated from a dispersed duplication event. Similarly, four of the *A. tenebrosa* SA8 genes originated from dispersed duplication events. However, there was no evidence of tandem duplication of SA8 genes in this species, with the two clustered SA8 genes arising from proximal duplication. Instead a genome doubling event in the genus *Actinia* [100] may have resulted in the distribution of SA8 genes across double the number of chromosomes compared to *T. stephensoni*

**Fig. 3.**
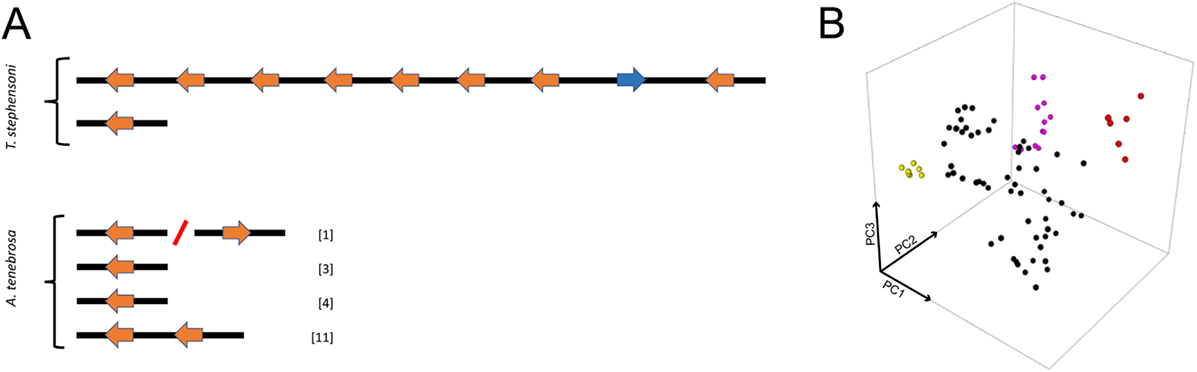
(A) Genomic arrangement and orientation of SA8 genes in *T. stephensoni* and *A. tenebrosa*. The inverted SA8 gene identified in the genome of *T. stephensoni* is depicted in blue. Numbers in square brackets indicate the scaffold number for *A. tenebrosa* SA8 genes. (B) Sequence space of the SA8 family. Sequence space of mature SA8 peptides reveals that they fall into four clusters, with the inverted SA8 *T. stephensoni* peptide found in the largest cluster (black)

Two separate instances of collinearity were detected between *T. stephensoni* and *A. tenebrosa* SA8 genes. The first of these collinear gene blocks included the *A. tenebrosa* SA8 gene and surrounding genes on scaffold 4 and the tandemly clustered *T. stephensoni* SA8 genes. The second SA8 gene on scaffold 1 of *A. tenebrosa* and the non-clustered *T. stephensoni* SA8 gene were the second instance of SA8 and other homologous genes evolving in a conserved order across both species. This indicates that collinearity can be observed in a subset of SA8 loci and suggests whole genome and/or segmental duplication has played a role in the evolution of the SA8 gene family in *A. tenebrosa*, with genes located on double the number of chromosomes.

Consistent with a recent expansion in *T. stephonsoni*, a high degree of sequence similarity and conserved gene structure present can be observed in *T. stephensoni*, but not *A. tenebrosa*, SA8 genes. While SA8 genes are comprised of 3–5 microexons in both species, high levels of variability are observed in the second exon in *A. tenebrosa* only. In *T. stephensoni*, there is high sequence similarity between the clustered SA8 genes, with the greatest sequence divergence associated with the non-clustered SA8 gene. Additionally, sequence space analysis revealed that SA8 peptides cluster into four groups across sea anemones (**Figure 3B**). SA8 peptides from *A. tenebrosa* were found in all four groups while most SA8 peptides from *T. stephensoni* (including the inverted venom SA8 peptide) were confined to a single group. The clustering patterns observed for *A. tenebrosa* and *T. stephensoni* sequences are reinforced in other species of Actinioidea and Metridioidea. Actinioidea species *Anthopleura dowii*, *Anthopleura buddemeieri*, *Stichodactyla haddoni* and *Aulactinia veratra* have SA8 peptides distributed across all four clusters. Conversely, SA8 peptides from Metridioidea species *E. diaphana*, *Nemanthus annamensis* and *Calliactus polypus* are restricted to one cluster, as are SA8 peptides identified in *Edwardsia carnea* and *N. vectensis* of superfamily Edwardsioidea. Therefore, despite the high copy numbers of SA8 genes in *T. stephensoni*, levels of sequence variation are lower in Metridioidea relative to Actinioidea. This finding favours the hypothesis that clustered SA8 genes observed in the *T. stephensoni* genome can be explained by a recent tandem duplication event and provides support for the hypothesis that tandem duplication may have played a key role in expansion of the SA8 family across the superfamily Metridioidea; however, further investigation using other actiniarian genomes will be needed to verify this.

Gene duplication has been shown to play a major role in the expansion of protein families in most venomous phyla. The snake venom metalloproteinase (SVMP) family in rattlesnakes originated from a single ancestral gene via a series of gene duplication events, resulting in 31 tandem SVMP genes in *Crotalus atrox* [103]. Similarly, venom phospholipase A_2_ (PLA_2_) genes cluster together in the genomes of multiple rattlesnake species, however, the number of genes in the cluster differs among *C. scutulatus*, *C. atrox*, and *C. adamanteus* [104]. A comparable pattern may emerge as SA8 genes are annotated in the genome of other Metridioidea species. As with SA8 clustered genes in *T. stephensoni*, the exons of clustered venom PLA_2_ genes in rattlesnakes are characterised by a conserved length and low levels of intron sequence variability [104]. Furthermore, tandem duplications are responsible for the evolution of defensin-like peptides in platypus venom [105–108]. Within order Actiniaria, a cluster of twelve genes encoding the Nv_1_ toxin were identified in the genome of *N. vectensis* [109]. These clustered Nv_1_ genes were proposed to have evolved via a birth and death mechanism [109, 110], a pattern consistent with the levels of nucleotide diversity found in the *T. stephensoni* SA8 genes. Furthermore, expansion events were reported to be more common in actiniarian-specific gene families – such as SA8 – compared to shared gene families [68], and tandem duplication was recently implicated in the evolution of the actiniarian ‘dominant toxin’ families [111]. Thus, there is a strong precedence for the expansion of the toxin gene families via duplication — both within sea anemones and other venomous lineages — and our data support the involvement of duplication events in the expansion and evolution of the SA8 family in *T. stephensoni*.

### SA8 peptides are associated with one of two expression patterns in *T. stephensoni*

The tissue distribution patterns of SA8 genes in *T. stephensoni* and *A. tenebrosa* were examined using several approaches. Differential gene expression analysis utilising previously generated tissue-specific reads for three envenomating structures of *A. tenebrosa* and three functional regions of *T. stephensoni* revealed distinct expression patterns of SA8 genes in each species. Only four of the six *A. tenebrosa* SA8 genes were differentially expressed (FDR ≤ 0.05, 4-fold) across the structures examined. Of these four differentially expressed *A. tenebrosa* SA8 genes (**Figure 4A**), two were upregulated in the acrorhagi (SA8_scaffold1_1 and SA8_scaffold3), one was upregulated in the mesenterial filaments (SA8_scaffold11_1), and one was upregulated in the tentacles (SA8_scaffold1_2). Conversely, in *T. stephensoni*, the nine transcripts corresponding to the 10 SA8 genes are differentially expressed across the tentacles (club-tips and tentacles), gastrodermis (actinopharynx and mesenterial filaments) and epidermis (body column and pedal disc) (**Figure 4B**). The majority of *T. stephensoni* SA8 genes are upregulated in the epidermis (four transcripts, five genes; SA8 clustered genes 1, 2, 6, 7 and 8) or the tentacles (three transcripts/genes; SA8 clustered genes 3, 4 and 5), although the non-clustered SA8 gene is upregulated in both the epidermis and tentacles.

**Fig. 4.**
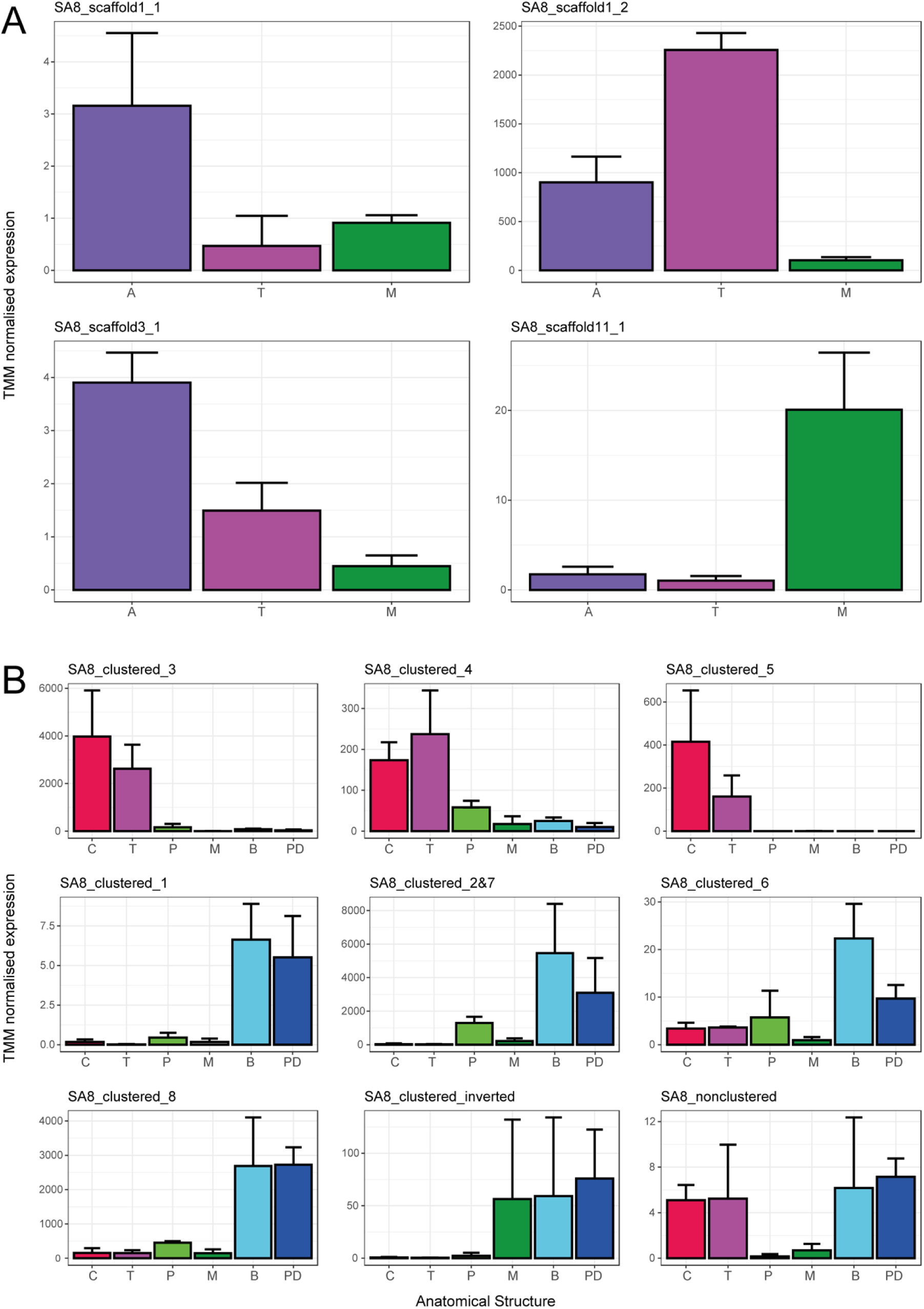
Expression of SA8 genes across functional regions in *A. tenebrosa* (A) and *T. stephensoni* (B). Each graph shows the tissue-specific expression pattern of a transcript that corresponds to a SA8 gene.

One transcript corresponds to two SA8 genes (SA8_clustered_2&7) in *T. stephensoni*. Bars represent the mean TMM-normalised expression value of the SA8 transcript in each anatomical structure, with error bars representing the standard deviation. Abbreviations: A= acrorhagi; C= club-tips; T= tentacle; P=actinopharynx; M= mesenterial filaments; B= body column; PD= pedal disc

Co-expression toxin module membership of all transcripts with significant BLASTp matches to SA8 peptides was assessed using weighted gene co-expression network analysis. In *T. stephensoni*, 11 transcripts (from eight genes) of the 15 transcripts (from ten SA8 genes) are assigned to two modules, each with a characteristic tissue-specific expression pattern (**Figure 5**); the turquoise module is characterised by upregulated expression in the tentacle tissue while the brown module is composed of putative toxins upregulated in column and pedal disc tissue. In *A. tenebrosa* SA8 transcripts are found in all four of the network modules identified (blue, brown, turquoise and yellow). Thus, the co-expression toxin modules associated with SA8 are defined by a specific expression pattern, and SA8 transcripts from *A. tenebrosa* are characterised by a greater range of expression profiles than those from *T. stephensoni*.

**Fig. 5.**
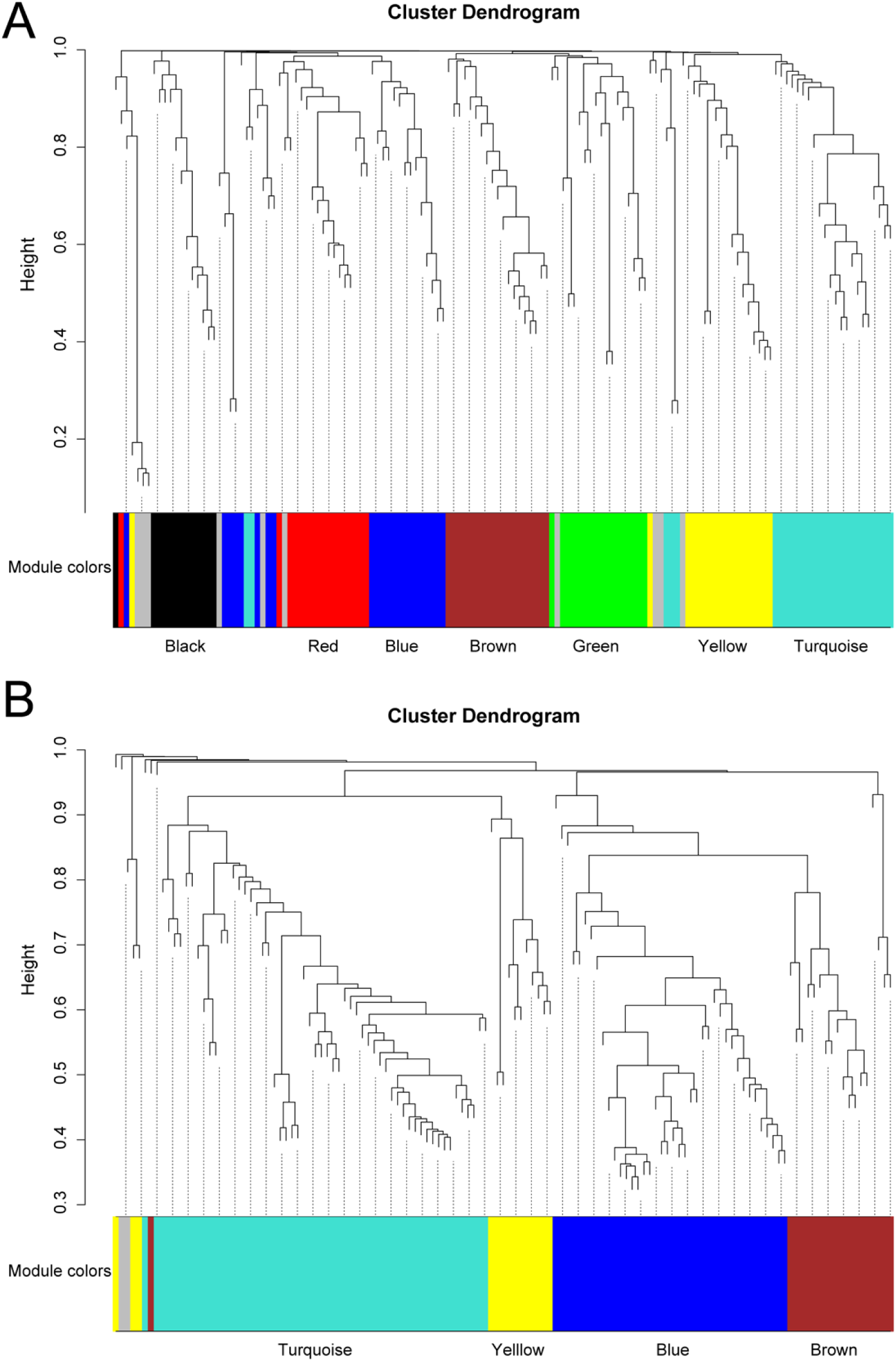
Cluster dendrogram of putative toxin transcripts from *T. stephensoni* (A) and *A. tenebrosa* (B) constructed using WGCNA. Each color represents one module. (A) Most *T. stephensoni* SA8 sequences were assigned to the turquoise and brown modules: turquoise = higher expression in the tentacles; brown = higher expression in the epidermis. The inverted *T. stephensoni* SA8 sequence was assigned to the black module, which was not associated with a specific expression pattern. Other modules: blue = higher expression in the mesenterial filaments; green= higher expression in the epidermis and tentacles; red= higher expression in the actinopharynx; yellow= higher expression in the tentacles and mesenterial filaments. (B) *A. tenebrosa* SA8 sequences were assigned to all four modules: blue = higher expression in the acrorhagi; brown= higher expression in the acrorhagi and tentacles; yellow= higher expression in the tentacles; turquoise = no distinct expression pattern. Putative toxin transcripts not assigned to a module are shown in grey

Following gene duplication, duplicates can be translocated, pseudogenised or retained, with expression divergence often occurring in those duplicates that are retained [110, 112–114]. One of the two major models to explain different functional roles of duplicated genes is subfunctionalisation, where the roles of the ancestral gene are partitioned between the duplicates [112, 115]. In venomous snakes, subfunctionalisation coincides with changes in the tissue distribution of toxin duplicates derived from salivary proteins [116]. By examining the tissue distribution of duplicated toxin gene families across venomous and non-venomous lineages, it was demonstrated that while toxin duplicates are restricted to the venom gland, their non-toxic counterparts continue to be expressed in the salivary glands of non-venomous reptiles [116]. Likewise, in the current study, we identified two major tissue-specific expression patterns for the SA8 family of *T. stephensoni*. Furthermore, evidence of subfunctionalisation was observed in *A. tenebrosa* with distinct tissue-specific expression patterns of duplicate SA8 genes, with subfunctionalisation of globin and toxin genes previously documented in this species [68, 117].

### A gene inversion event leads to recruitment of a SA8 peptide into the venom of *T. stephensoni*

An inversion event was also identified within the tandemly duplicated SA8 gene cluster of *T. stephensoni*. This inverted SA8 gene shows substantial sequence divergence from other clustered SA8 genes, with only the non-clustered *T. stephensoni* SA8 gene having greater sequence divergence. Like other *T. stephensoni* SA8 genes, the inverted SA8 gene is upregulated in the epidermis; however, it is unique in that elevated expression can also be observed in gastrodermis structures, specifically the mesenterial filaments. Mass spectrometry imaging confirmed that the peptide corresponding to the calculated mass of the *T. stephensoni* SA8 venom peptide is located in the same anatomical structures (**Figure 6**). Furthermore, the transcript corresponding to the inverted SA8 gene is found in a separate co-expression module (black) to other *T. stephensoni* SA8 peptides.

**Fig. 6.**
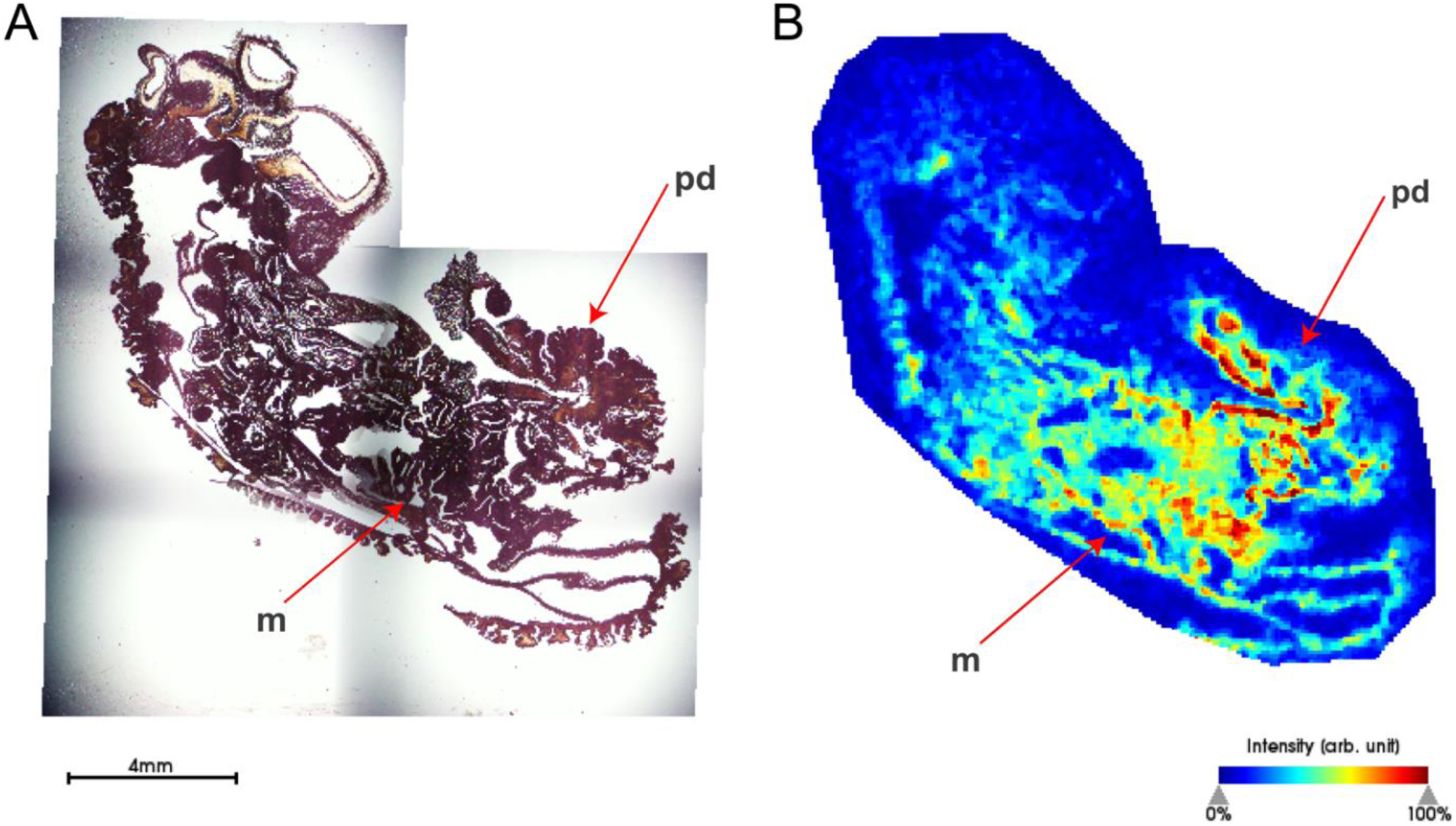
Peptide corresponding to calculated mass of the *T. stephensoni* SA8 venom peptide is abundant in the gastrodermis and epidermis (A) H&E stain of *T. stephensoni* section. (B) putative peptide observed at 5409 *m/z* with greatest abundance in the mesenterial filaments (m) and pedal disc (pd)

In addition, two peptides from the milked venom proteome with similarity to previously identified *Anemonia viridis* SA8 peptides were mapped back to the inverted SA8 gene. These peptides represent alternate splice isoforms of the inverted SA8 gene, with different transcriptional start sites that maintain the open reading frame and have identical mature peptides. As none of the peptides identified in the milked venom of *A. tenebrosa* had significant similarity to SA8 sequences, the SA8 sequences in this species are probably non-venom peptides. This suggests that the inversion event, in concert with tandem duplication, may have facilitated the recruitment of this SA8 peptide into the venom of *T. stephensoni*, with SA8 not found in the venom of any other actiniarian species. Furthermore, the inverted gene is found in a well-supported clade on a branch sister to a Metridioidea specific clade (**Figure 7**), indicating that the recruitment of this orthologue into the venom may have only occurred in the *Telmatactis* or related acontiarian sea anemones.

**Fig. 7.**
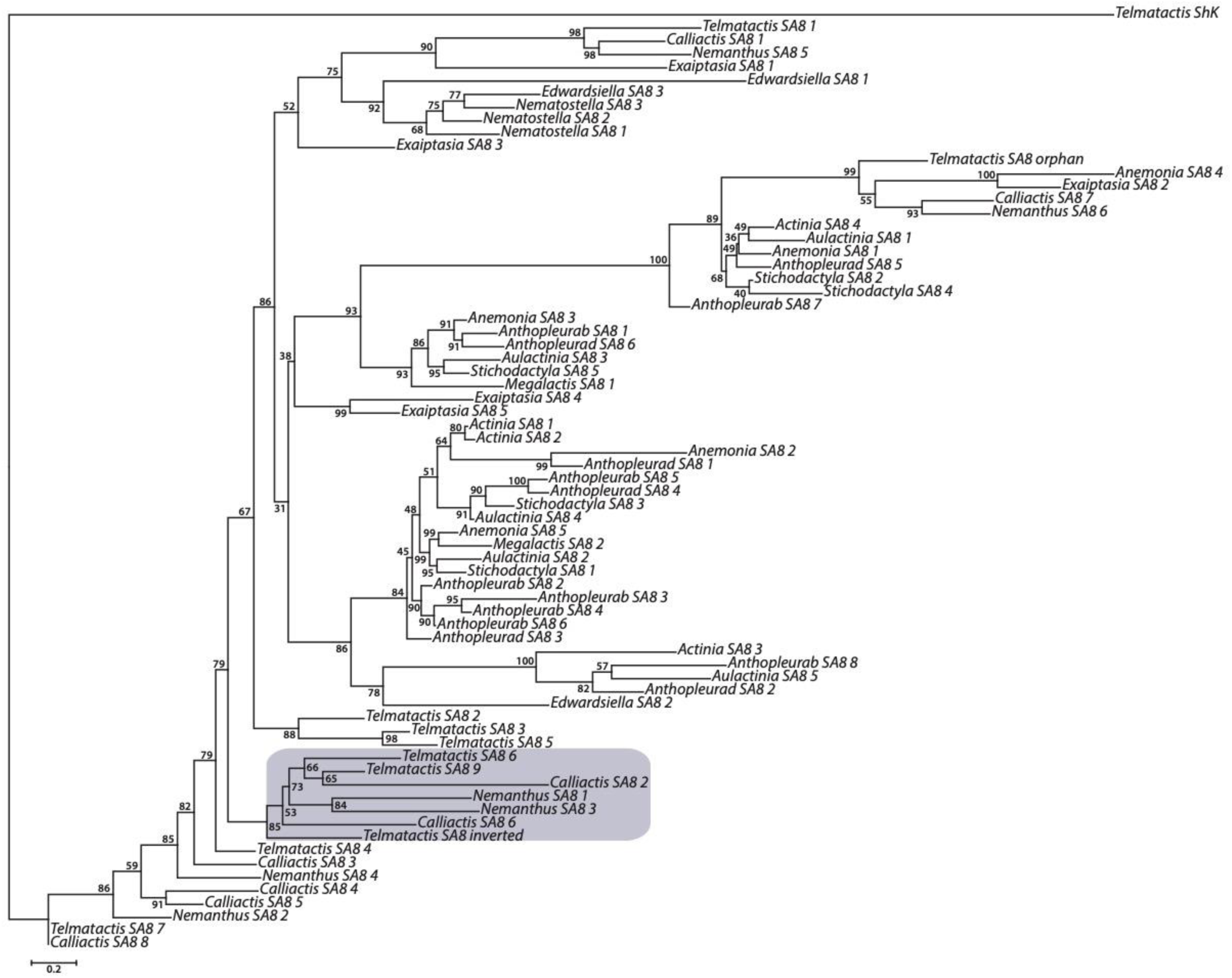
Phylogenetic relationships among 71 sea anemone sequences from the SA8 gene family were inferred under maximum likelihood in IQ-TREE. The inverted SA8 gene from *T. stephensoni* is found in a well-supported clade on a branch sister to a Metridioidea-specific clade

Neofunctionalisation occurs when the duplicated gene assumes a role distinct from that of the original gene, and it is the second major model used to account for functional divergence following gene duplication [114, 115]. Although considered a rare event [116], neofunctionalisation has occurred in multiple snake venom gene families [104, 118–122]. For example, in the snake-venom PLA_2_ gene family, duplication and subsequent inversion of a basic PLA_2_ gene resulted in the first acidic-PLA_2_ gene, and thus this combination of events can represent a scenario that enables neofunctionalisation [104]. We propose an inversion event and subsequent escape from a gene regulatory network has allowed the venom *T. stephensoni* SA8 gene to undergo neofunctionalization and a novel gene expression program. This gene also appears to have had a high number of amino acid replacements compared to the other clustered SA8 genes in *T. stephensoni*.

### The SA8 family is distinct from the ShK and possesses a unique disulfide network

All SA8 putative toxins in *T. stephensoni* had a cysteine spacing consistent with motif 3, described by Kozlov and Grishin [29], which has previously been observed in SA8 putative toxins from *Anemonia viridis* (P0DMZ3, P0DMZ4, P0DMZ5, P0DMZ6, P0DMZ7), sea anemone type 1 potassium channel toxins with a ShKT domain (P29187, P29186, P81897, Q9TW91) [29] and cysteine-rich domain (CRD) of snake venom cysteine-rich secretory proteins (CRISPs) [123]. Sequence space analysis [27] revealed that ShK-like proteins, CRISPs and SA8 putative toxins are clearly delineated into three groups (**Figure 8A**), reinforcing the unique sequence composition of SA8 putative toxins and indicating that they represent a single distinct protein family.

**Fig. 8.**
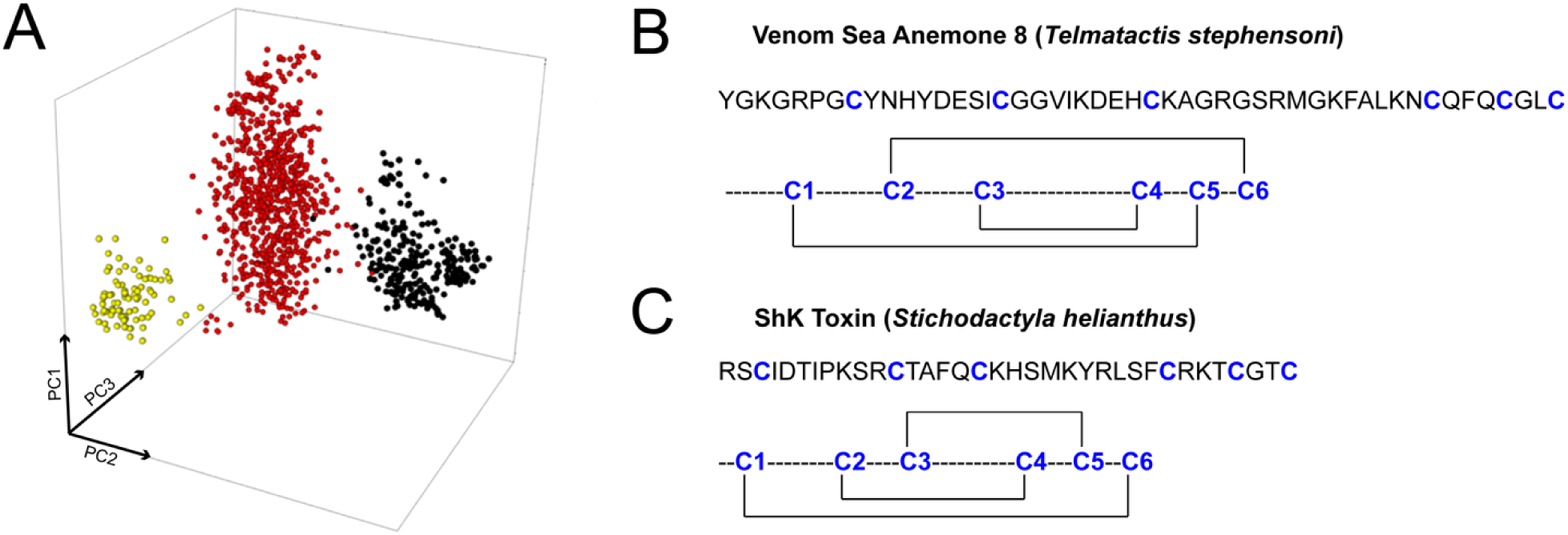
Sequence space of the SA8 family (A) and disulfide connectivities of venom SA8 (B) and ShK (C). (A) The SA8 family (yellow) forms a distinct cluster from the ShKT domains of ShK-like (red) and CRISP (black) peptides [27]. (B) The C1-C5, C2-C6, C3-C4 disulfide connectivity of the mature venom SA8 peptide from *T. stephensoni*. (C) The C1-C6, C2-C4, C3-C5 disulfide connectivity of ShK toxin from *S. helianthus*

All SA8 peptides are characterised by a conserved glycine residue and FA dyad (**Supplementary Figure 1A**). This dyad has not been reported in other toxin groups and further research is required to determine its functional significance. Few SA8 peptides contain the KY dyad characteristic of ShK, although most contain a KY dyad alternative (QY, KW, TY, RF); however, the two protein families possess distinct sequence patterns and exon-structure. SA8 peptides contain more residues between the pro-cleavage site and final cysteine, resulting in larger proteins, but only ShK-like sequences have an extension at the C terminus. SA8 genes are consistently composed of 3–5 microexons (**Supplementary Figure 1B**), while ShK genes are composed of 2–3 exons. Additionally, large introns are present in all SA8 genes, but particularly so in *A. tenebrosa*, with a 17-kb intron identified in this species. Therefore, although ShK-like and SA8 peptides have a similar cysteine spacing pattern, these two families are resolved into discrete groups when examining their other characteristics.

To examine the structure of the SA8 peptide present in milked venom and determine if it is similar to that of ShK, we expressed this venom peptide and characterised it using LC-MS/MS and NMR spectroscopy. The His6-MBP-SA8 fusion protein was the dominant protein expressed in transformed *E. coli* cells following IPTG induction. SA8 was liberated from the 50.5 kDa fusion protein using TEV protease (**Supplementary Figure 2**), then purified by affinity chromatography followed by RP-HPLC (**Supplementary Figure 3A**). The LC-MS profile of the fraction corresponding to SA8 showed a pure peak whose molecular mass (MH^+^ *m/z*= 5438 Da) was in good agreement with the expected mass (theoretical MH^+^ *m/z*= 5438.23 Da) (**Supplementary Figure 3B**). The recombinant SA8 co-elutes on RP-HPLC with native SA8, with only a very slight difference in elution time (11.44 min and 11.38 min, respectively) due to the presence of an additional C-terminal glycine on the recombinant peptide (**Supplementary Figure 4**). Furthermore, the observed mass of the recombinant SA8 was 6 Da less than that of the reduced form of the peptide (MH^+^ *m/z*= 5444.23 Da), indicating formation of three disulfide bonds. LC-MS/MS analysis of pepsin-digested SA8 revealed the disulfide-bond connectivity to be C1-C5, C2-C6, and C3-C4 (**Figure 8B, Supplementary Table 3**). This pattern differs from the ShK disulfide framework (C1-C6, C2-C4, C3-C5) (**Figure 8B**), supporting the hypothesis that SA8 is not a simple extension of the ShK fold.

A 1D ^1^H NMR spectrum of recombinant SA8 acquired at pH 3.5 and 25°C revealed poor dispersion of amide-proton resonances region (**Supplementary Figure 5**). Subsequently, several 1D spectra were recorded at temperatures over the range 5–30 °C in 5 °C increments and pH values 2–7 (**Supplementary Figure 6 and 7**), but no significant improvement in peak dispersion was observed. Moreover, a sample of the recombinant peptide that was reduced and unfolded, then oxidatively refolded *in vitro* gave the same spectral dispersion as the recombinant peptide, indicating that we are dealing with the most stable fold of this sequence (**Supplementary Figure 8**). Thus, the NMR data indicate that the *T. stephensoni* SA8 venom peptide adopts a partially disordered structure in solution, unlike ShK which has a well-defined structure [18]. Overall, we report the SA8 family is associated with unique disulfide-bond connectivities, exon-intron structure, and sequence spaces that distinguish it from ShK.

### Functional activity of SA8 peptides remains unknown

ShK is a 35-residue peptide isolated from the sea anemone *S. helianthus* [14] that potently inhibits K_V_1 channels through a conserved KY functional dyad [124–126]. The presence of KY dyad alternatives in multiple SA8 sequences suggests this putative toxin family may have some activity against K^+^ channels [127]. First, we tested the effects of 100 nM recombinant SA8 on the following human K_V_ channels: K_V_1.1, K_V_1.2, and K_V_1.3. Whole-cell K_V_1.2 currents were recorded sequentially on the same transiently transfected CHO cell, before (control trace, black) and after perfusion of the cell with 100 nM SA8 (red trace) (**Figure 9A**). At equilibrium block, SA8 caused ∼10% reduction in current amplitude. This weak block was almost fully reversible upon washing the perfusion chamber with inhibitor-free solution. Both the association and dissociation rate of SA8 were rapid; development of equilibrium block and recovery up to ∼85% of control current took ∼1 min. In contrast, 100 nM SA8 did not inhibit K_V_1.1 or K_V_1.3 (**Supplementary Figure 9A and B**). We performed a concentration-response experiment for the inhibition of K_V_1.2 channels by SA8 (**Figure 9B**) and determined the reciprocal of the slope of the fitted line yielded an IC_50_ of 40.8 ± 4.9 µM (n=4).

**Fig. 9.**
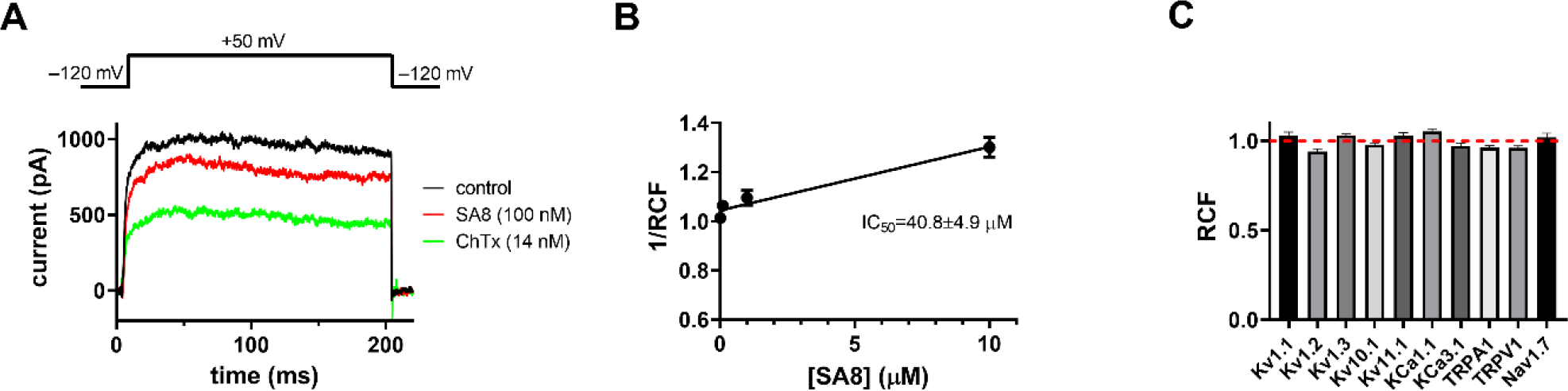
SA8-induced inhibition of hK_V_1.2 and selectivity profile of SA8. (A) Representative whole-cell current traces were recorded for hK_V_1.2 using the voltage protocol shown above the raw current traces every 15 s in the absence (black, control) and presence of 100 nM SA8 (red) and 14 nM charybdotoxin (ChTx, green) as positive control. (B) Low affinity, concentration-dependent block of hK_V_1.2 channels by SA8 was determined fitting a straight line to the reciprocal of the remaining current fraction (1/RCF) plotted as a function of SA8 concentration. Remaining current fraction (RCF) was calculated as *I/I_0_*, where *I_0_* is the peak current in the absence and *I* is the peak current at equilibrium block in the presence of SA8 at concentrations of 0.01, 0.1, 1, and 10 µM (filled circles). Points on the linear concentration-response curve represent the mean of four independent measurements where the error bars represent SEM. The line was drawn using linear least squares fit and the reciprocal of the slope of the best fit yielded an IC_50_ of 40.8 ± 4.9 µM. (C) The effect of SA8 (100nM, except for K_V_10.1 which was tested at 1µM) on the peak currents was reported as the remaining current fraction (RCF). Bars represent the mean of 4–6 independent measurements; error bars indicate the SEM. Data are shown for the following channels: Hk_V_1.1, hK_V_1.2, hK_V_1.3, hK_V_10.1, hK_V_11.1, mK_Ca_1.1, hKCa3.1, hTRPA1, hTRPV1, and hNa_V_1.7 (for details of the expression systems, solutions, and voltage protocols, see Materials and Methods, and for raw current traces see Supplementary Figure 9). SA8 did not inhibit either of the investigated channels at the applied concentrations

To reveal the selectivity profile of SA8 we assayed its effect on other K^+^ channels as well. We tested its activity on K_V_10.1 and K_V_11.1 due to their slightly different structure compared to K_V_1.x channels [128–130]. We also tested the effect of SA8 on K_Ca_1.1, the large conductance voltage- and Ca^2+^-activated channel, and K_Ca_3.1, a Ca^2+^-activated K^+^ channel. Moreover, as SA8 is suggested to be a defensive peptide and possibly involved in painful stings, channels involved in pain recognition were also tested including the voltage-gated sodium channel Na_V_1.7, and the transient receptor potential channels TRPA1 and TRPV1. We used positive controls (2 µM astemizole for K_V_10.1, 10 mM TEA^+^ for K_Ca_1.1, 20 nM TRAM-34 for K_Ca_3.1, 50 µm HC-030031 for TRPA1, and 50 µm capsazepine for TRPV1) diluted freshly in ECS in order to confirm ion channel expression and proper operation of the perfusion system. We found that most of the channels shown in **Figure 9** **and Supplementary Figure 9** are not inhibited significantly by SA8 at the indicated concentrations except for the modest block of the hKv1.2 current.

Injection of *D. melanogaster* with recombinant SA8 at a dose of 0.5 µg per fly did not result in paralysis or death within 24 h. In contrast, neurotoxic spider-venom peptides with insecticidal activity cause paralysis and death at dose of 0.005 µg per fly in *D. melanogaster* [99]. Thus, the recombinant SA8 peptide may not exert any toxic effects in insects and only causes a weak inhibition of human Kv1.2 channels. However, the presence of this peptide in the venom indicates that it probably has at least an auxiliary function in venom and the tissue distribution of the inverted SA8 peptide suggests that this putative peptide toxin may target the neuronal channels of vertebrate predators in order to fulfil the ecological function of predator deterrence [58]. Further functional testing is required to determine the biological activity of this venom.

## Conclusions

Based on sequence, structural and genomic data, we find that SA8 putative toxins are a distinct family that has only superficial similarities to ShK. Thus, characterising the SA8 family offers an opportunity to explore a novel group of putative toxins that may possess pharmacological properties that make them useful as pharmacological tools or therapeutic leads.

We report that the SA8 family is expanded in *T. stephensoni* but not *A. tenebrosa* relative to other actiniarian species. The two genomic locations of SA8 genes in *T. stephensoni* form two distinct collinear blocks with two of the four loci of SA8 genes in *A. tenebrosa*, indicating that segmental duplication may have played a role in evolution of the SA8 family. However, the expansion of clustered SA8 genes in *T. stephensoni* appears to be the consequence of proximal and tandem duplication events.

While members of this family are expressed in the neuronal cells of *N. vectensis*, a SA8 peptide has been recruited to the venom of *T. stephensoni* following an inversion event. This inverted SA8 gene is distinct from other members of the SA8 family across multiple levels and represents a potential neofunctionalisation event in the SA8 gene family. In contrast, the absence of non-inverted SA8 peptides from the venom of *T. stephensoni* and *A. tenebrosa*, as well as the previously reported neuronal localisation of SA8-like peptides suggests that other SA8 genes may encode non-venom neuropeptides.

Furthermore, while we have not yet determined the functional activity of venom SA8 peptides, we have shown that they are structurally distinct from ShK toxins and possess a unique disulfide bond pattern. We propose that the inverted *T. stephensoni* SA8 peptide has a role in defence against predators, as either a toxin or an auxiliary venom protein, based on its distribution across the body plan.

Additional testing will be required to determine the exact function of the inverted SA8 and the SA8 family in general, but given the findings of the current study, the genetics origins and diversification of this putative toxin family should be investigated in other sea anemones. Future research should determine whether the patterns we report are conserved across Actinioidea and Metridioidea, particularly whether venom SA8 peptides are only found when there is an inversion within a cluster of tandemly duplicated genes.

## Statements and Declarations

## Acknowledgements

The authors thank the QUT Marine group for their help and advice caring for animals, and acknowledge the QUT Molecular Genetics Research facility (MGRF) for use of their facilities. The authors acknowledge the facilities and scientific and technical assistance of the Australian Microscopy & Microanalysis Research Facility at the Centre for Microscopy and Microanalysis, The University of Queensland. Computational resources and services used in this work were provided by the High Performance Computing and Research Support Group, Queensland University of Technology, Brisbane, Australia. We would also like to thank for Ira Cooke for his expert advice.

## Author Contributions

Conceptualization, L.M.A., K.A.E., R.S.N. and P.J.P.; Transcriptomic data analysis, L.M.A.; Genomic data analysis, L.M.A., Z.K.S., C.A.V.B., H.L.S., J.M.S., V.L.C., M.J.P., K.J.D. and P.J.P.; sequence space analysis: T.M.A.S.; Structural and functional analysis: K.A.E., M.U.N., T.G.S., S.G. and D.C.C.W.; Proteomic data analysis, E.A.B.U., B.M. and B.R.H.; Original draft preparation, L.M.A., K.A.E., Z.K.S., T.G.S., R.S.N. and P.J.P; Draft review and editing, L.M.A., K.A.E., Z.K.S., T.M.A.S., M.U.N., T.G.S., C.A.V.B., H.L.S., J.M.S., E.A.B.U., B.M., B.R.H., S.G., D.C.C.W., V.L.C., M.J.P., K.J.D., D.A.H., G.P., G.F.K., A.P., R.S.N. and P.J.P.; Supervision, E.A.B.U., D.C.C.W., D.A.H., G.P. G.F.K., A.P., R.S.N. and P.J.P. All authors have read and agreed to the published version of the manuscript.

## Data Accessibility

Raw RNA reads for *T. stephensoni* and *A. tenebrosa* are available at the NCBI SRA under BioProjects PRJNA728752, PRJNA350366 and PRJNA313244. Tissue-specific *T. stephensoni* reads (PRJNA728752) were used for differential gene expression analysis and transcriptome-based gene model prediction. Tissue-specific reads for *A. tenebrosa* (PRJNA350366) were used for differential gene expression analysis [33], while whole-organism reads (PRJNA313244) were used for transcriptome-based gene model prediction. Accession numbers for each tissue sample are given in the table below. Assembled genomes for *T. stephensoni* and *A. tenebrosa* have been uploaded to NCBI genome and are available via accession numbers JALIZF000000000 and JALIZE000000000, respectively. The MS proteomics data have been uploaded to the ProteomeXchange Consortium via the PRIDE [131] partner repository with the dataset identifier PXD029717.

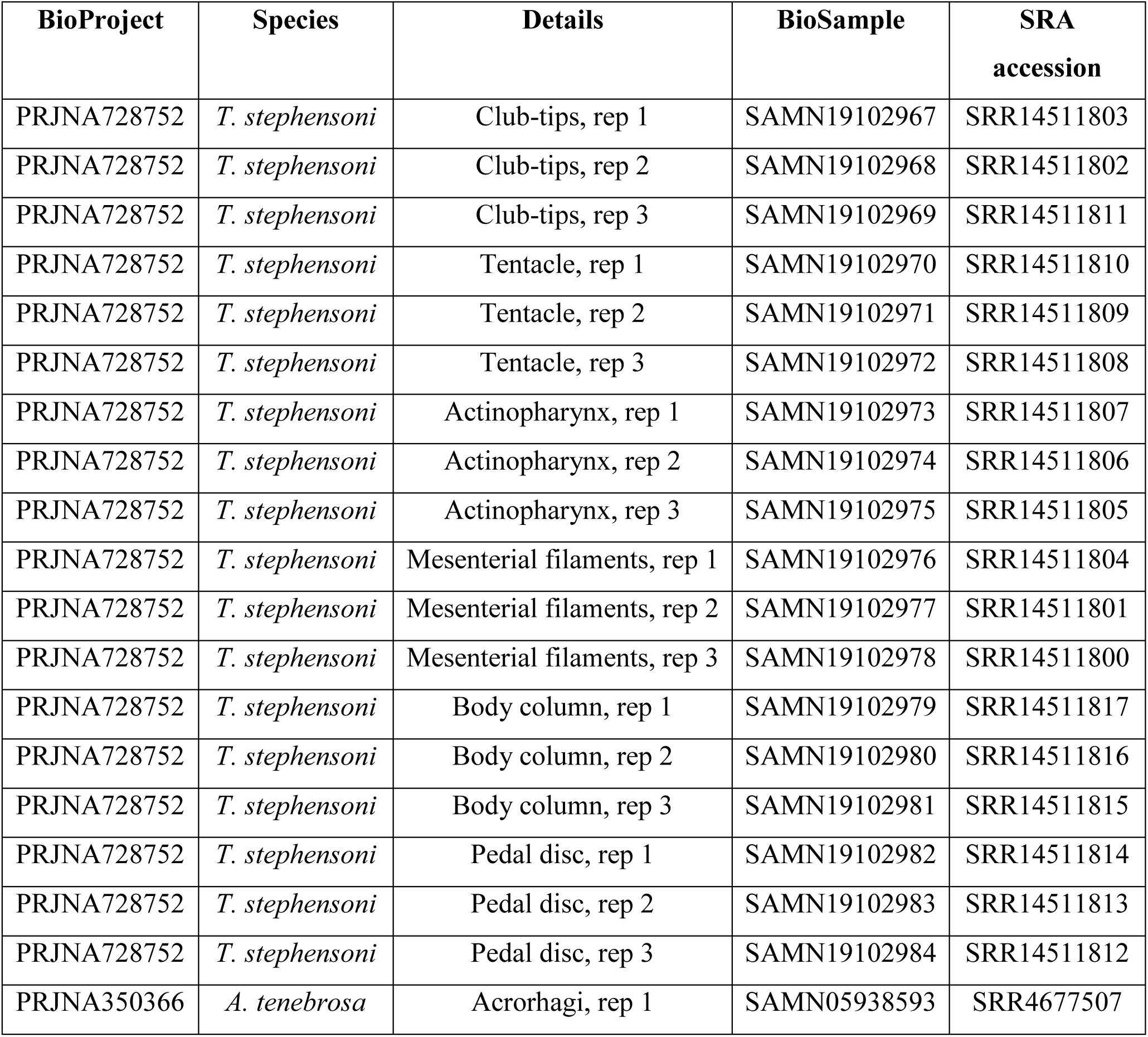
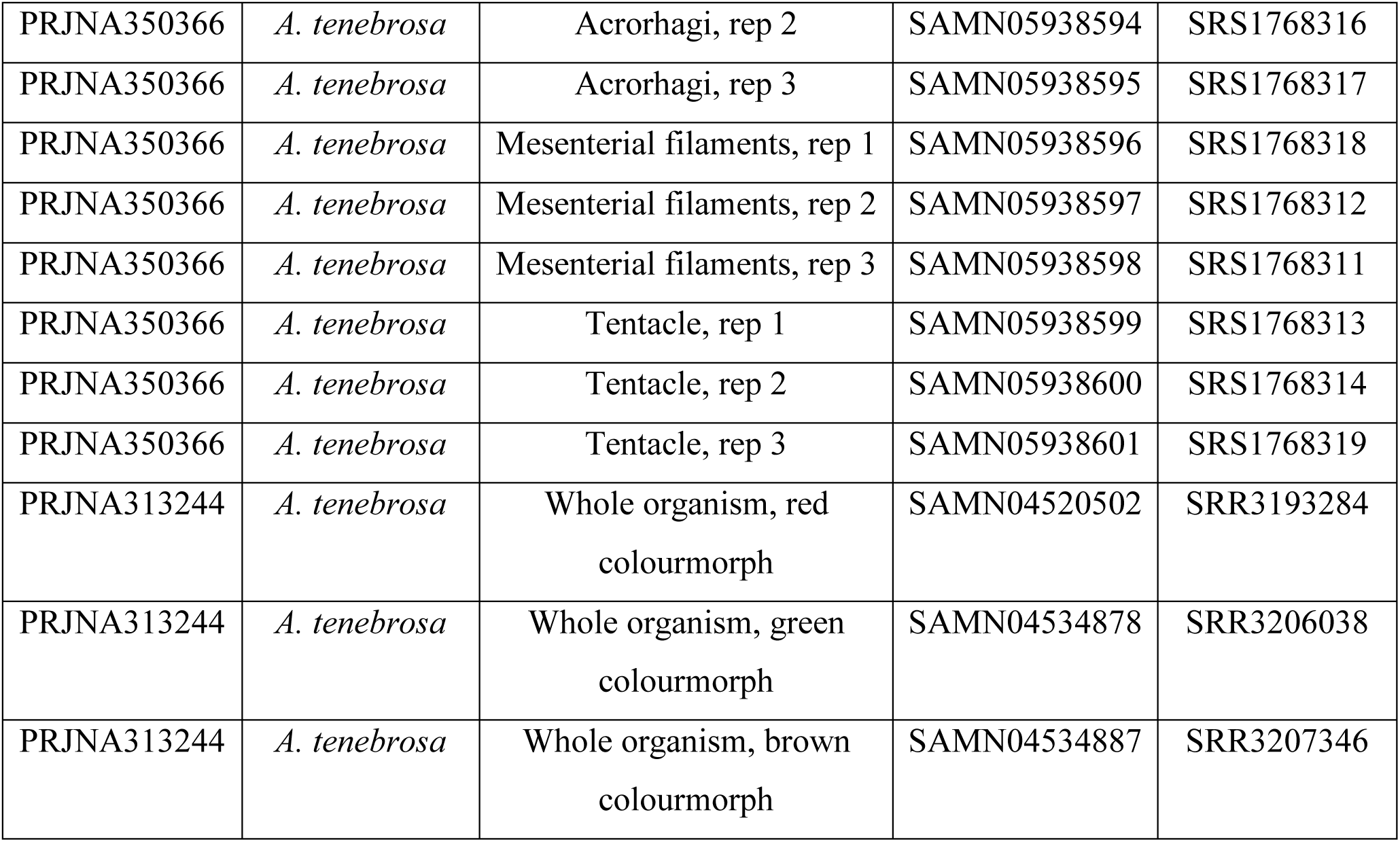

| BioProject | Species | Details | BioSample | SRA<br>accession |
| --- | --- | --- | --- | --- |
| PRJNA728752 | *T. stephensoni* | Club-tips, rep 1 | SAMN19102967 | SRR14511803 |
| PRJNA728752 | *T. stephensoni* | Club-tips, rep 2 | SAMN19102968 | SRR14511802 |
| PRJNA728752 | *T. stephensoni* | Club-tips, rep 3 | SAMN19102969 | SRR14511811 |
| PRJNA728752 | *T. stephensoni* | Tentacle, rep 1 | SAMN19102970 | SRR14511810 |
| PRJNA728752 | *T. stephensoni* | Tentacle, rep 2 | SAMN19102971 | SRR14511809 |
| PRJNA728752 | *T. stephensoni* | Tentacle, rep 3 | SAMN19102972 | SRR14511808 |
| PRJNA728752 | *T. stephensoni* | Actinopharynx, rep 1 | SAMN19102973 | SRR14511807 |
| PRJNA728752 | *T. stephensoni* | Actinopharynx, rep 2 | SAMN19102974 | SRR14511806 |
| PRJNA728752 | *T. stephensoni* | Actinopharynx, rep 3 | SAMN19102975 | SRR14511805 |
| PRJNA728752 | *T. stephensoni* | Mesenterial filaments, rep 1 | SAMN19102976 | SRR14511804 |
| PRJNA728752 | *T. stephensoni* | Mesenterial filaments, rep 2 | SAMN19102977 | SRR14511801 |
| PRJNA728752 | *T. stephensoni* | Mesenterial filaments, rep 3 | SAMN19102978 | SRR14511800 |
| PRJNA728752 | *T. stephensoni* | Body column, rep 1 | SAMN19102979 | SRR14511817 |
| PRJNA728752 | *T. stephensoni* | Body column, rep 2 | SAMN19102980 | SRR14511816 |
| PRJNA728752 | *T. stephensoni* | Body column, rep 3 | SAMN19102981 | SRR14511815 |
| PRJNA728752 | *T. stephensoni* | Pedal disc, rep 1 | SAMN19102982 | SRR14511814 |
| PRJNA728752 | *T. stephensoni* | Pedal disc, rep 2 | SAMN19102983 | SRR14511813 |
| PRJNA728752 | *T. stephensoni* | Pedal disc, rep 3 | SAMN19102984 | SRR14511812 |
| PRJNA350366 | *A. tenebrosa* | Acrorhagi, rep 1 | SAMN05938593 | SRR4677507 |
| PRJNA350366 | *A. tenebrosa* | Acrorhagi, rep 2 | SAMN05938594 | SRS1768316 |
| PRJNA350366 | *A. tenebrosa* | Acrorhagi, rep 3 | SAMN05938595 | SRS1768317 |
| PRJNA350366 | *A. tenebrosa* | Mesenterial filaments, rep 1 | SAMN05938596 | SRS1768318 |
| PRJNA350366 | *A. tenebrosa* | Mesenterial filaments, rep 2 | SAMN05938597 | SRS1768312 |
| PRJNA350366 | *A. tenebrosa* | Mesenterial filaments, rep 3 | SAMN05938598 | SRS1768311 |
| PRJNA350366 | *A. tenebrosa* | Tentacle, rep 1 | SAMN05938599 | SRS1768313 |
| PRJNA350366 | *A. tenebrosa* | Tentacle, rep 2 | SAMN05938600 | SRS1768314 |
| PRJNA350366 | *A. tenebrosa* | Tentacle, rep 3 | SAMN05938601 | SRS1768319 |
| PRJNA313244 | *A. tenebrosa* | Whole organism, red<br>colourmorph | SAMN04520502 | SRR3193284 |
| PRJNA313244 | *A. tenebrosa* | Whole organism, green<br>colourmorph | SAMN04534878 | SRR3206038 |
| PRJNA313244 | *A. tenebrosa* | Whole organism, brown<br>colourmorph | SAMN04534887 | SRR3207346 |

## Ethics Approval

This study was confirmed as Outside the Scope and exempt from the need for University Animal Ethics Committee (UAEC) review, approval, and monitoring in conformity with the Australian code for the care and use of animals for scientific purposes.

