## Supplemental Materials for "Genomic, Functional and Structural Analyses Reveal Mechanisms of Evolutionary Innovation within the Sea Anemone 8 Toxin Family"

**Supplementary Table 1** Repeats masked in the genomes of *A. tenebrosa* and *T. stephensoni*

| Repeat Class | ***A. tenebrosa*** | | ***T. stephensoni*** | |
|---|---|---|---|---|
| | **Nt masked** | **% masked** | **Nt masked** | **% masked** |
| **DNA** | 9,829,854 | 3.14 | 7,692,305 | 1.72 |
| **LINE** | 4,695,670 | 1.50 | 7,554,675 | 1.69 |
| **LTR** | 41,149,192 | 13.16 | 30,839,449 | 6.89 |
| **MITE** | 26,200,481 | 8.38 | 41,367,077 | 9.24 |
| **RC** | 417,313 | 0.13 | 1,917,466 | 0.43 |
| **Satellite** | 112,468 | 0.04 | 1,040,798 | 0.23 |
| **Simple repeat** | 204,204 | 0.07 | 324,491 | 0.07 |
| **SINE** | 1,540,055 | 0.49 | 1,456,064 | 0.33 |
| **SINE-like** | 98,150 | 0.03 | 54,894 | 0.01 |
| **Unknown** | 16,659,416 | 5.33 | 29,977,185 | 6.70 |

**Supplementary Table 2** CAFE was used to identify expanded gene families encoding putative toxins (fixed Orthogroups) across actiniarian superfamilies Actinioidea, Edwardsioidea and Metridioidea. Species: Telmatactis = *T. stephensoni*; Paraphelliactis = *P. xishaensis*; Exaiptasia = *E. diaphana*; Actinia = *A. tenebrosa*; Nematostella = *N. vectensis*. Uncharacterised toxin gene families have been designated U# or Z#

| Species | Expanded Orthogroup | Putative Toxin Gene Families |
|---|---|---|
| Telmatactis | OG0000200 | FactorV-like_L1 |
| Telmatactis | OG0000228 | Z3 |
| Telmatactis | OG0000508 | Z6 |
| Telmatactis | OG0000570 | FactorV-like_L1 |
| Telmatactis | OG0000636 | CREC |
| Telmatactis | OG0000700 | FactorV-like_L3 |
| Telmatactis | OG0000841 | IGFBP-like_L2 |
| Telmatactis | OG0000924 | Peptidase_M12A_L2 |
| Telmatactis | OG0001131 | FactorV-like_L3 |
| Telmatactis | OG0001408 | Peptidase_M12A_L7 |
| Telmatactis | OG0001452 | Peptidase_M12A_L1, Peptidase_M12A_L4 |
| Telmatactis | OG0001503 | FactorV-like_L1 |
| Telmatactis | OG0001654 | Peptidase_S1_L1 |
| Telmatactis | OG0001663 | PLA2_L1 |
| Telmatactis | OG0001684 | ShK-like_L2 |
| Telmatactis | OG0001912 | FactorV-like_L3 |
| Telmatactis | OG0002282 | IGFBP-like_L3 |
| Telmatactis | OG0007619 | Sea_Anemone_8 |

| | | |
|---|---|---|
| Telmatactis | OG0008704 | ShK-like_L2 |
| Telmatactis | OG0016521 | FactorV-like_L3 |
| Telmatactis | OG0018001 | U15 |
| Paraphelliactis | OG0000215 | IGFBP-like_L1, IGFBP-like_L2, IGFBP-like_L3 |
| Paraphelliactis | OG0000450 | IGFBP-like_L2, IGFBP-like_L3 |
| Paraphelliactis | OG0000778 | IGFBP-like_L3 |
| Paraphelliactis | OG0001257 | FactorV-like_L1, FactorV-like_L2, FactorV-like_L3 |
| Paraphelliactis | OG0001433 | Peptidase_S1_L1 |
| Paraphelliactis | OG0001607 | FactorV-like_L3 |
| Paraphelliactis | OG0001625 | Kazal-like_L2 |
| Paraphelliactis | OG0003153 | FactorV-like_L2 |
| Paraphelliactis | OG0010929 | Peptidase_S1_L1 |
| Paraphelliactis | OG0023536 | IGFBP-like_L3 |
| Exaiptasia | OG0000017 | IGFBP-like_L3 |
| Exaiptasia | OG0000152 | FactorV-like_L1 |
| Exaiptasia | OG0000159 | FactorV-like_L1 |
| Exaiptasia | OG0000200 | FactorV-like_L1 |
| Exaiptasia | OG0000233 | IGFBP-like_L3 |
| Exaiptasia | OG0000369 | FactorV-like_L2 |
| Exaiptasia | OG0000921 | IGFBP-like_L3 |
| Exaiptasia | OG0001029 | Peptidase_M12A_L1, Peptidase_M12A_L6 |
| Exaiptasia | OG0001066 | Peptidase_M12A_L1 |
| Exaiptasia | OG0001194 | Peptidase_M12A_L1 |

| | | |
|---|---|---|
| Exaiptasia | OG0001344 | Peptidase_M12A_L5 |
| Exaiptasia | OG0001452 | Peptidase_M12A_L1, Peptidase_M12A_L4 |
| Exaiptasia | OG0001625 | Kazal-like_L2 |
| Exaiptasia | OG0002274 | IGFBP-like_L3 |
| Exaiptasia | OG0006279 | IGFBP-like_L3 |
| Actinia | OG0000203 | IGFBP-like_L2, IGFBP-like_L3 |
| Actinia | OG0002170 | U11_L1 |
| Actinia | OG0002840 | IGFBP-like_L3 |
| Actinia | OG0006503 | FactorV-like_L2 |
| Nematostella | OG0010929 | Peptidase_S1_L1 |
| Nematostella | OG0023536 | IGFBP-like_L3 |

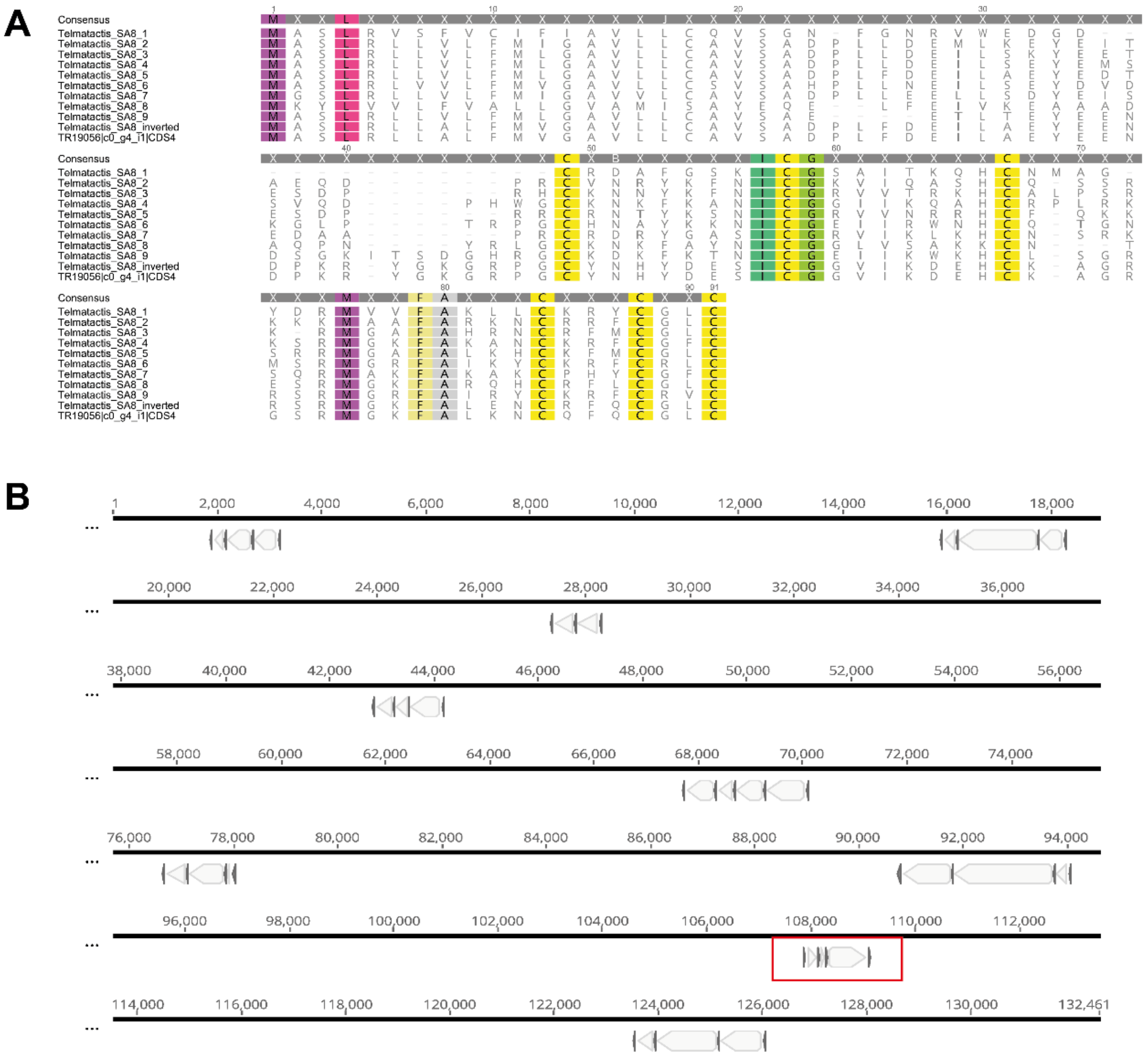


**Supplementary Fig. 1** Sequence and structure of *T. stephensoni* SA8 sequences. (A) Alignment of SA8 gene and peptide sequences from *T. stephensoni*, with conserved cysteine framework, glycine residue, and FA dyad highlighted. (B) The nine clustered genes of *T. stephensoni* are composed of 3–5 microexons and large introns. The inverted *T. stephensoni* SA8 gene is indicated in red

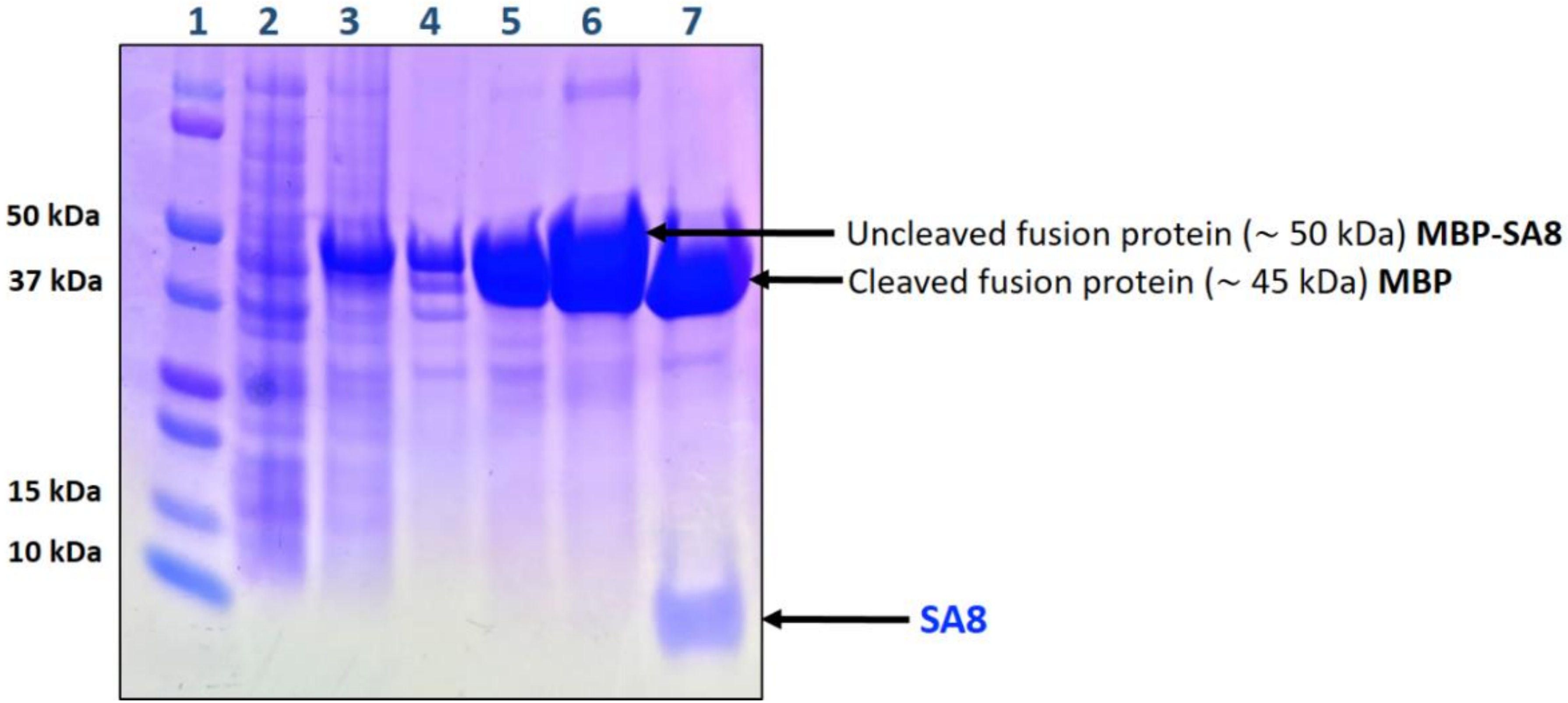


**Supplementary Fig. 2** SDS-PAGE gel stained with Coomassie Blue showing samples of the His6-MBP-SA8 fusion protein obtained during different steps of expression and purification. Lane 1: molecular mass standards; Lanes 2 and 3: *E. coli* cells pre- and post-induction with IPTG; Lane 4: sucrose extract; Lane 5: periplasmic extract (5 mM $MgCl_2$); Lanes 6 and 7: purified MBP fusion protein before and after cleavage with TEV protease

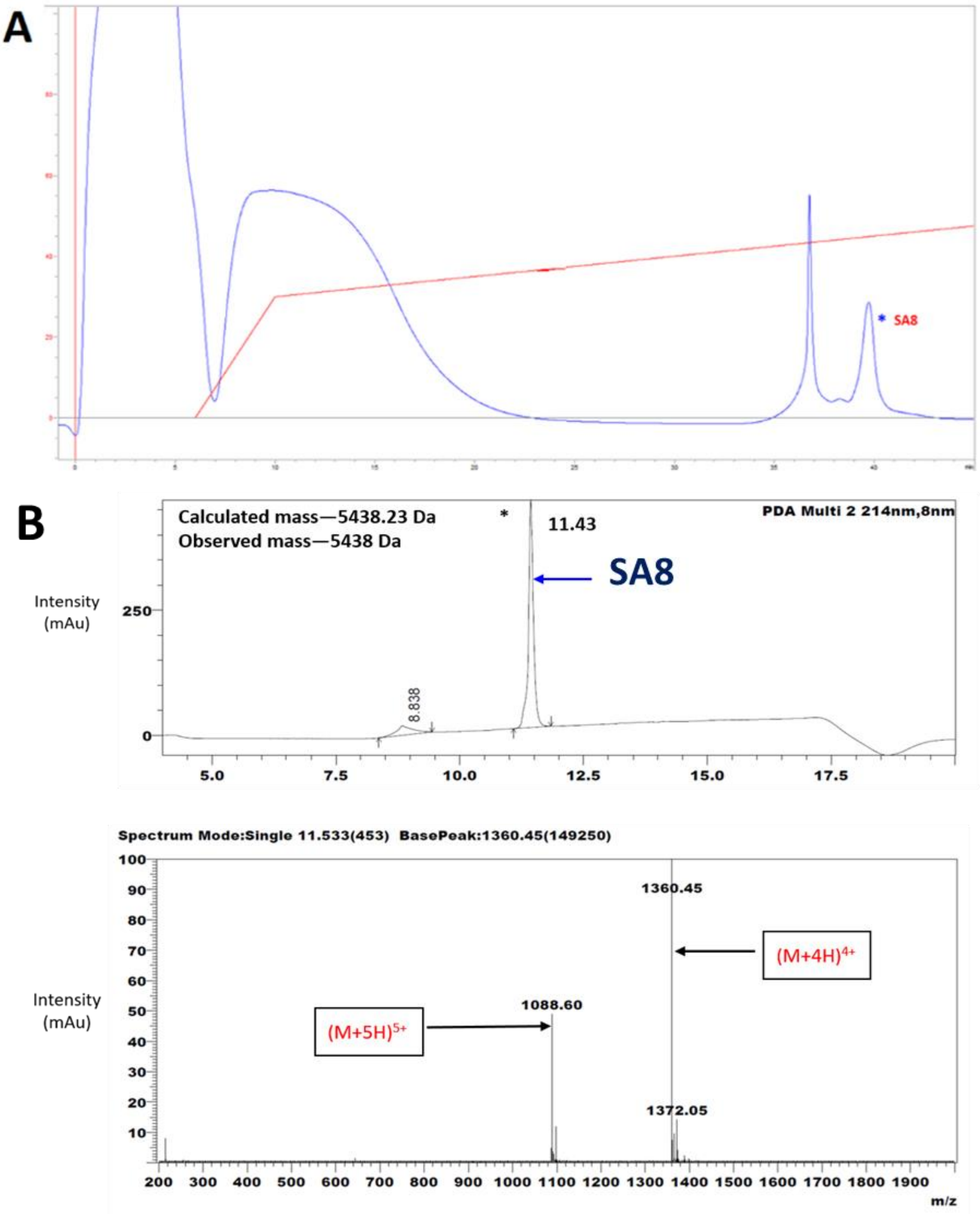


**Supplementary Fig. 3** (A) RP-HPLC purification of recombinant SA8. The peptide was separated on a Vydac C18 column (10 x 250 mm) using a flow rate of 1 mL/min and a 40 min linear gradient of 30–50% solvent B (0.1% TFA in acetonitrile), as indicated by the red line. (B) LC-MS profile of oxidised SA8 after purification using RP- HPLC. The observed mass is consistent with formation of three disulfide bonds

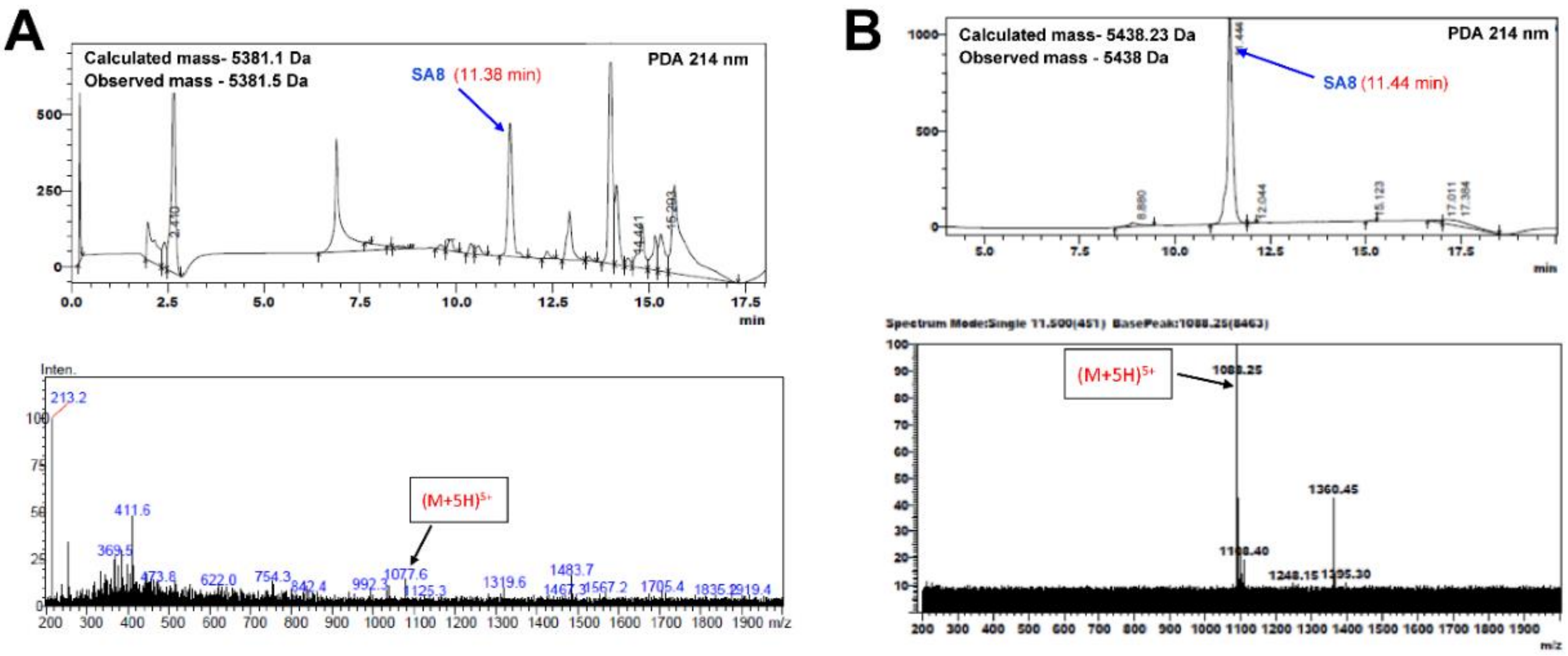


**Supplementary Fig. 4** Comparison of LC-MS profiles for co-eluted native and recombinant SA8. (A) LC-MS profile of the native SA8 peptide. (B) LC-MS profile of the recombinant SA8 peptide

**Supplementary Table 3** Direct mass spectrometric characterisation of disulfide linkages in the recombinant SA8 peptide. The main diagnostic pepsin fragments appeared as MS2 precursors with high intensity

| Disulfide connectivity | Corresponding pepsin fragment |
|---|---|
| C1–C5 | GYGKGRPGCY—QCGL |
| C2–C6 | DESICGGVI—C |
| C4–C3 | KNCQF–KDEHCKA |
| C1–C5,C6–C2 | GYGKGRPGCY–QCGLC–CGGVI |

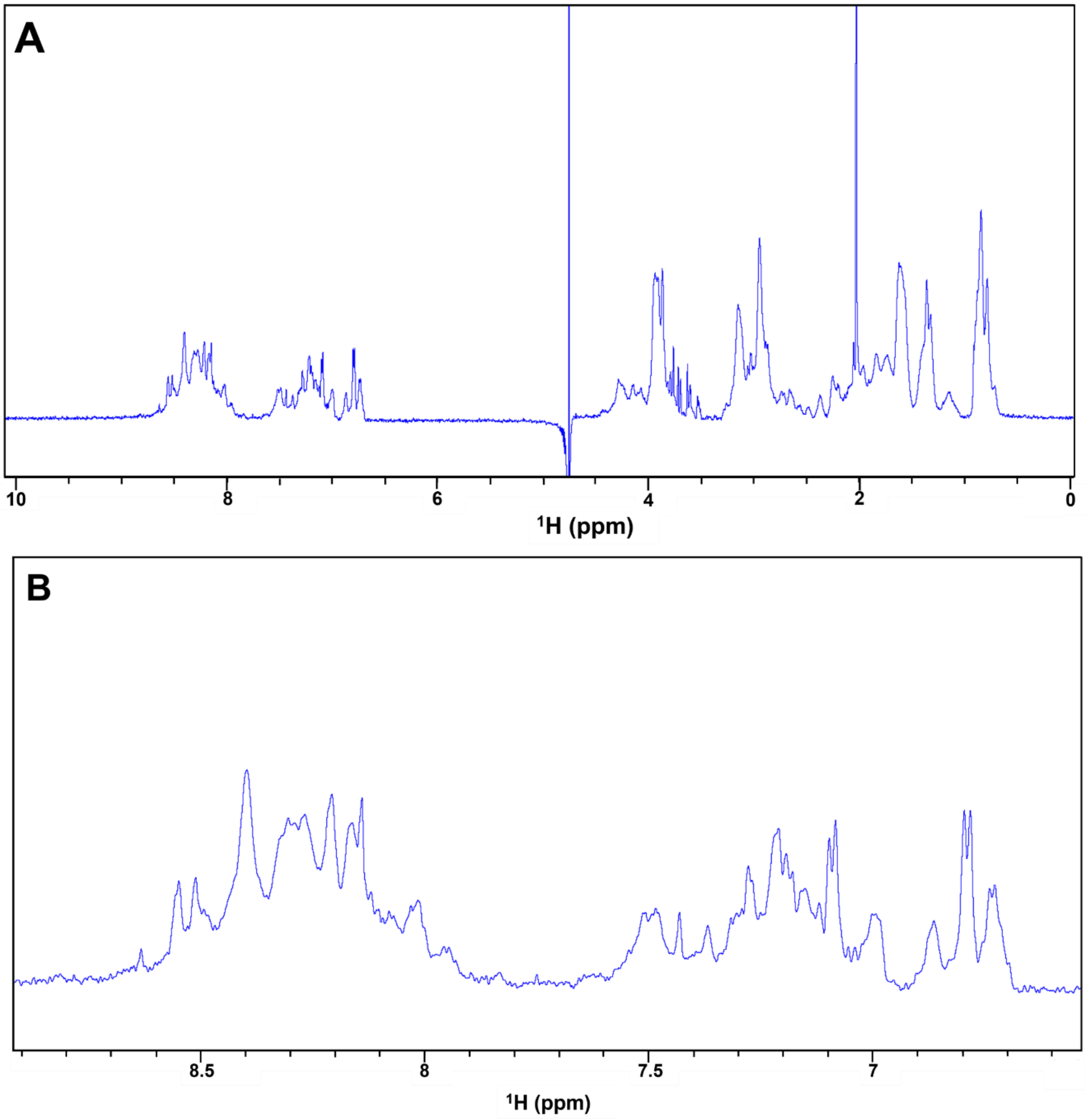


**Supplementary Fig. 5** (A) Full 1D $^{1}$H NMR spectrum of recombinant SA8 recorded on Bruker 600 MHz NMR spectrometer at pH 3.5 and 298 K and 256 scans. (B) Expanded amide/aromatic region

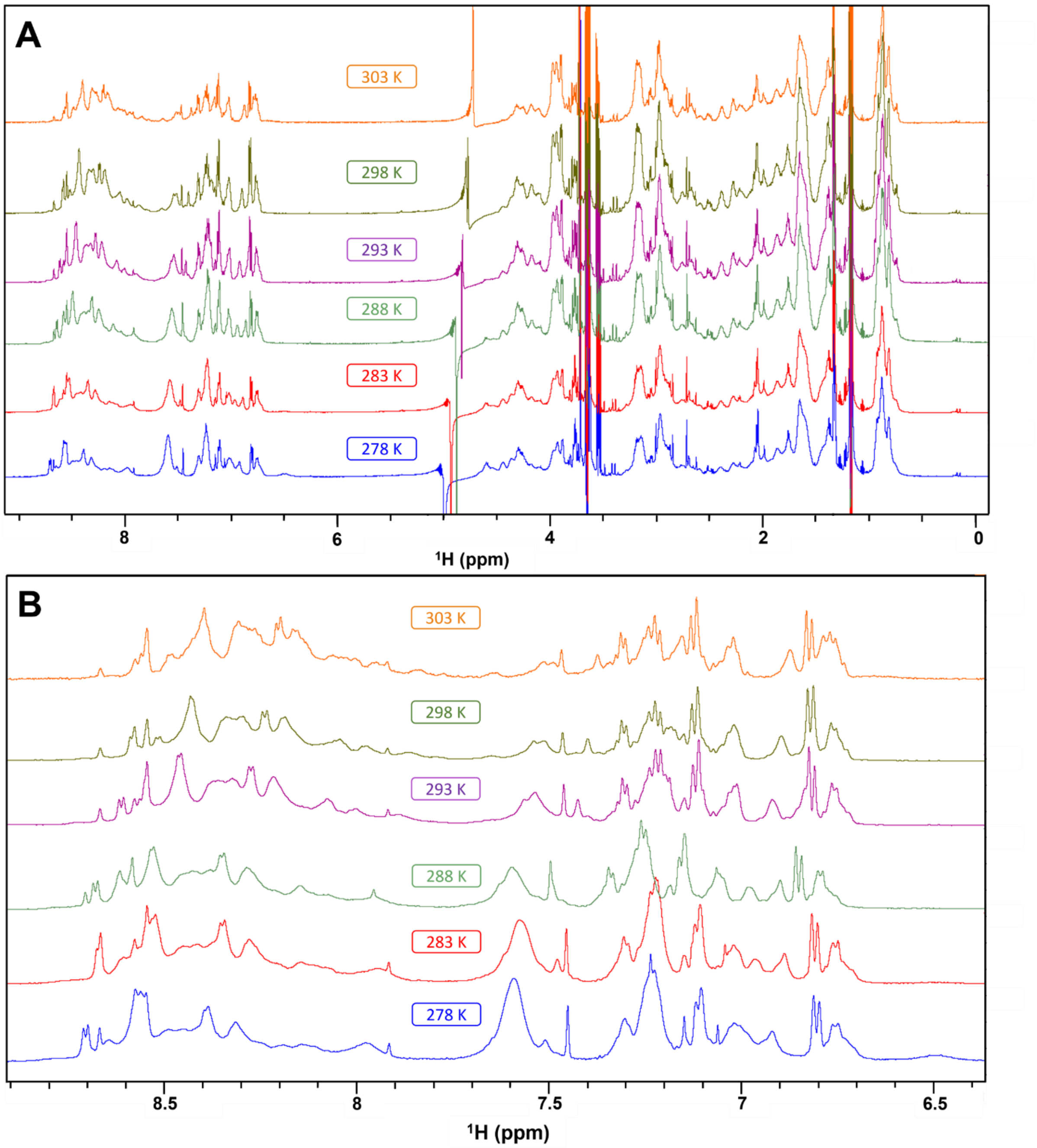


**Supplementary Fig. 6** (A) 1D $^{1}$H NMR spectra of SA8 recorded on Bruker 600 MHz NMR spectrometer. Spectra were acquired at pH 3.5 and different temperatures over the range 5–30 °C. (B) Expanded amide/aromatic region

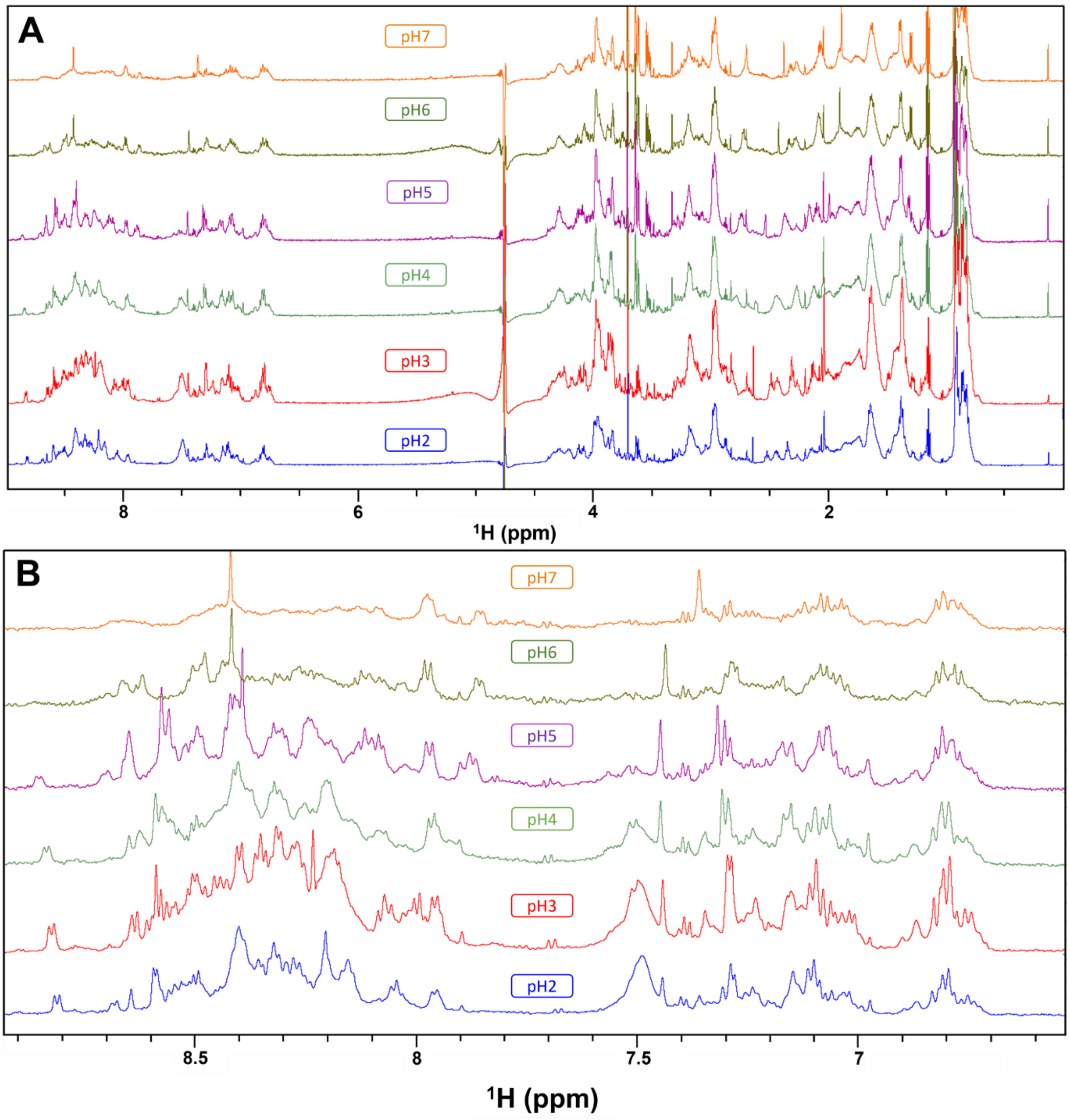


**Supplementary Fig. 7** (A) 1D $^{1}$H NMR spectra of SA8 recorded on Bruker 600 MHz NMR spectrometer. Spectra were acquired at 298 K using 256 scans, over the pH range 2-7. (B) Expanded amide/aromatic region

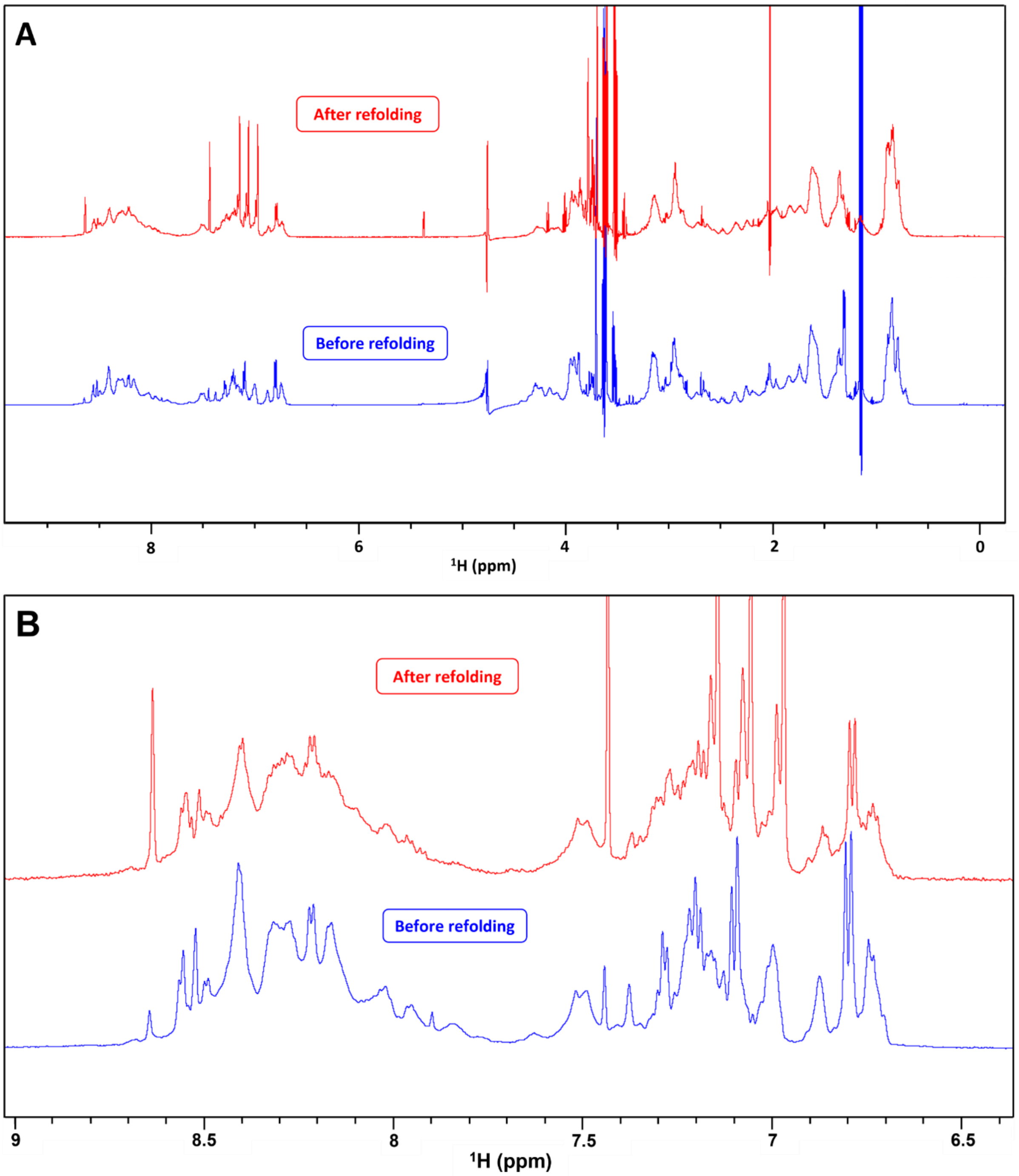


**Supplementary Fig. 8** (A) 1D $^1$H NMR spectra of SA8 recorded on Bruker 600 MHz NMR spectrometer. Spectra were acquired at pH 3.5 and 298 K using 256 scans. (B) Expanded amide/aromatic region. N.B**.** The sharp strong peaks in the spectra are from low molecular weight molecules

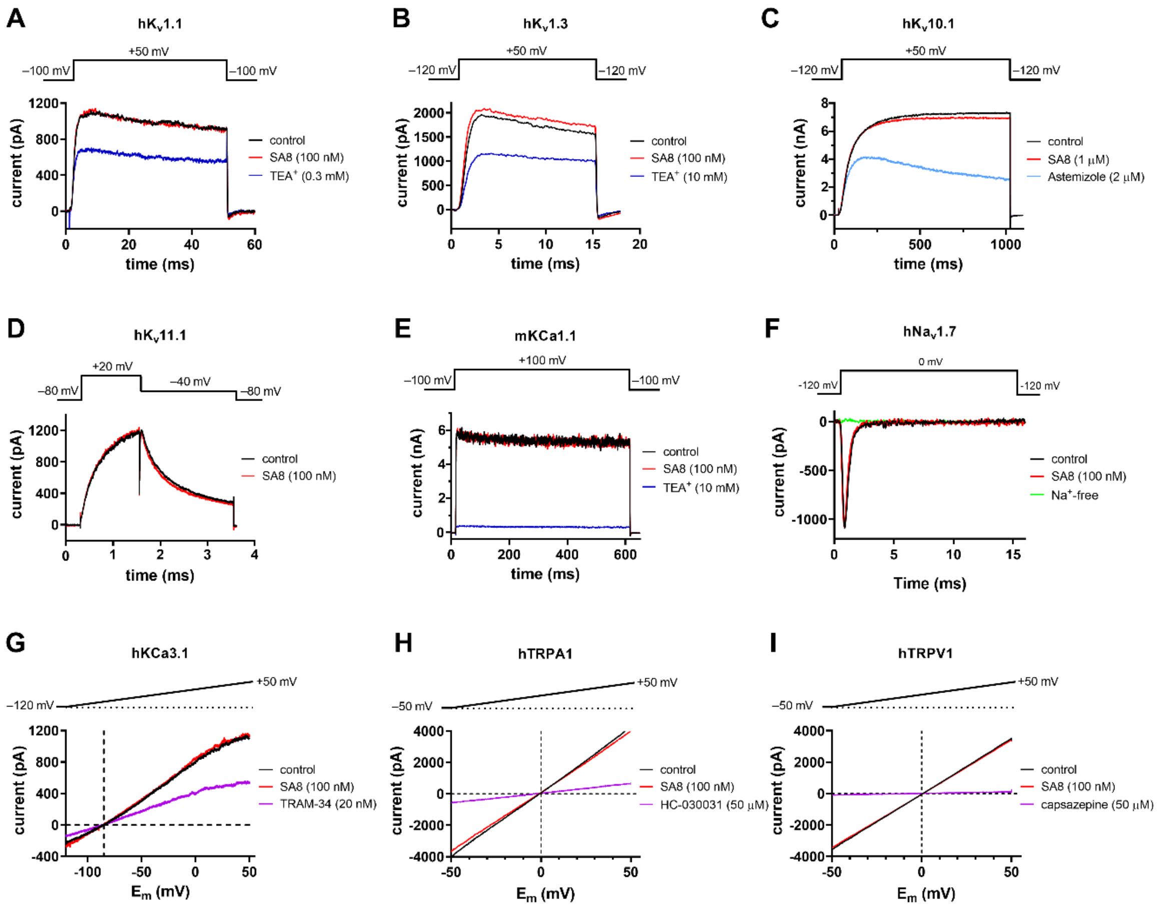


**Supplementary Fig. 9** SA8 has no effect on several voltage-gated and $Ca^{2+}$-activated ion channels. Current traces recorded before application of SA8 (control, black), after 1-2 min perfusion with SA8 (red, at 100 nM SA8 concentration except panel C where the SA8 concentration was 1 μM), and after perfusing the recording chamber with control solutions ($Na^+$-free ECS solution or specific inhibitors of the ion channel, colored traces). Data are shown for the following channels: $Hk_V1.1$ (A), $hK_V1.3$ (B), $hK_V10.1$ (C), $hK_V11.1$ (D), mKCa1.1 (E), $hNa_V1.7$ (F), hKCa3.1 (G), hTRPA1 (H), and hTRPV1 (I). For details on the expression systems, solutions, and voltage protocols, see Materials and Methods. For hKCa3.1, hTRPA1, and hTRPV1, the currents were recorded in response to a voltage ramp, corrected for ohmic leakage and then displayed as a function of test potential ($E_m$). The horizonal dashed line shows the zero current level and the vertical dashed line indicates the expected reversal potential for $K^+$ (based on the Nernst equation)
